# Development of iPSC-derived urothelial organoids towards investigating the effect of hormones on host-defense to urinary tract infections

**DOI:** 10.64898/2026.08.29.747866

**Authors:** Adam J Bindas, Zizhuang Fang, Jos Boekhorst, Aline Marie Fernandes, Jerry M Wells

## Abstract

Recurrent urinary tract infection represents a substantial unmet public health in women. Local administration of estradiol has been shown to reduce recurrence, however *in vitro* models of the female urinary tract remain limited and the mechanisms underlying the effects of estradiol are incompletely understood. Here, we describe a novel iPSC organoid differentiation protocol and its application to establish a multilayered Transwell barrier culture model. Estradiol treatment resulted in reduced expression of innate antimicrobial peptides and cytokines, together with increased expression of demannosylation pathways. To our knowledge, this is the first iPSC organoid-derived model of the urinary tract, which provides a platform for investigating interactions between the urothelium, urobiome and hormonal environment.

## Introduction

UTIs are one of the most common reasons to visit a general practitioner^1^, with almost half of women experiencing at least one UTI during their lifetime.^2^ Recurrence is common with, 20- 30% of women experiencing another infection within 6 months. Post-menopausal women have an increased risk of UTI including recurrent UTI (rUTI), suggesting that host factors may play a role in reinfection risk.^3,4^ Estrogen levels decline substantially after menopause^5,6^ and estrogen administration has been shown to modify urinary tract host defense.^7,8^ Meta-analysis of clinical data indicates that vaginal, but not oral estrogen therapy may reduce rUTI recurrence.^9^ While estrogen supplementation may be beneficial, it remains unclear how estrogen effects the urothelium and what role other hormones altered following menopause may play in urothelial host defense.

Testosterone^10,11^ levels are also diminished in post-menopausal populations. In addition, increased norepinephrine levels and sympathetic activity have been reported in postmenopausal women, and may partly reflect the loss of systemic estrogen.^12,13^ Sex hormones can modulate innate immune inflammation signaling providing a rationale for examining their effects in a urothelial model.^14^

While UTIs are much more common in women than men^2^, many existing urothelial cell lines including HBLAK, 5637, TERT-NHUC A, and SV-HUC-1 are isolated from male patients, which may limit their suitability for investigating the effects of female sex hormones. Of these cell lines, the spontaneously immortalized HBLAK line can form a stratified epithelium following exposure to patient urine^15^, however it contains genetic modifications consistent with early-stage tumors.^16^ Urine-derived stem cells, which can be obtained non-invasively from patient urine have been expanded into organoids.^17–19^ Although bladder cancer cells are known to regularly shed into urine, healthy patient urine likely contains a wide variety of cell lineages, including mesoderm-derived renal cells.

Primary or iPSC urothelial organoids provide an alternative approach to modeling non-malignant urothelium and have been demonstrated to recapitulate tissue-level epithelial diversity, while allowing investigation of donor-specific and genetically defined phenotypes^1^ Primary urothelial organoids have predominantly been generated from tumor tissue for studies on bladder cancer, although organoids derived from normal tissue have also been described ^20,21^ Qu et al. recently described a method to generate urothelial organoids from iPSCs, where they reported spontaneous stratification and differentiation of organoid cultures.^22^ However, a need exists to develop a robust approach for establishing organoid-derived, multilayered urothelial barrier cultures suitable for co-culture with bacteria.

Here, we investigate the response of female iPSC-derived urothelial organoids to sex hormones. To our knowledge, we report the first multilayered urothelial barrier culture derived from human iPSC-derived urothelial organoids. This model provides a platform for investigating host-microbe interactions and mechanisms relevant to rUTI.

## Materials and Methods

### iPSC cell culture

iPSC cells (NAS2 or EDi002-A) were thawed from cryopreserved stocks and cultured as previously described^23^, on 6-well plates (Stem Cell, 38015) with 2 mL of mTeSR medium (Stem Cell, 85857). Culture surfaces were coated with 10 µg/mL Vitronectin XF (Stem Cell, 07180) in CellAdhere buffer (Stem Cell, 7183). Cells were fed daily and passaged at 80% confluence at a 1:6 ratio onto freshly coated plates for further expansion or at a ratio of 1:30 onto 24-well plates for differentiation. A concentration of 0.5 µg/cm^2^ of iMatrix-511 recombinant laminin E8 fragment (AMS Bio, 892012) was used to coat the differentiation culture surface. For passaging, 1 mL of ReLeSR (Stem Cell, 100-0483) was after removal of mTeSR. When gaps began to form in the cell layer, 2 mL of mTeSR medium was added, and the remaining adherent cells were gently lifted. The cell suspension was transferred to a fresh culture surface and in 1 mL or 0.5 mL of medium per well in the 6- and 24-well plates, respectively.

### Definitive endoderm induction and urothelial cell differentiation

Once iPSC cultures achieved 80% confluence, definitive endoderm (DE) differentiation medium was introduced for 4 days, as previously described^24^. DE medium was consisted of RPMI-1640 with GlutaMAX supplement (Fisher, 61870044) containing 100 ng/mL of Activin A (Fisher, 17169311), 5 µM of CHIR99021 (Bio-Techne, 44231/10) and 1X B27 (Gibco, 17504-044). Next, urothelium (U) differentiation medium was supplied for an additional 10 days. U medium consisted of RPMI-1640 with GlutaMAX supplement (Fisher, 61870044) containing 10 µM all-trans retinoic acid, 100 ng/mL human FGF10 (Biolegend, 559304), and 1x KGM-Gold SingleQuots (Lonza, 192152). For the 4 final days, cultures were maintained in urothelial maturation (UM) medium, consisting of U medium supplemented with 1 µM of Troglitazone (Stem Cell, 73892) and 1 µM PD153035 (Sigma, SML0564).

At day 18 of differentiation, the cell cultures were dissociated using 1 mL of TrypLE Express (Gibco, 12604013) and incubated at 37°C until the cells detached from the surface (approx. 12 minutes). Cells were lightly lifted with a 1000 mL pipette before being transferred to a 15 mL tube and centrifuged (300G for 5 minutes at 4°C). Following supernatant removal, the pellet was resuspended in 300 µL of Matrigel (Corning, 734-1101) and seeded as small droplets (50 µl) into six wells of a 24-well plate. Cultures were incubated for at least 15 minutes at 37°C before the addition of organoid expansion medium supplemented with 10 µM of Y-27632 ROCK inhibitor (Tocris, Cat. No. 1254/10).

### Organoid culture

Expansion medium consisted of Advanced DMEM/F12 (Fisher, 11540446) with 50% conditioned L-WRN medium (ATCC, CRL-3276), 1X B27, 1X N-2 supplement (Gibco, 11520536), 1X HEPES (Gibco, 15630056), 1X GlutaMAX (Gibco, 35050061), 1.25 mM N-acetyl-cysteine (Sigma, A9165), 10 mM Nicotinamide (Tocris, 4106), 0.5 µM A 83-01 (Tocris, 2939), 50 ng/mL EGF (Stem Cell, 78006.1), 10 µM SB202190 (Stem Cell, 17100102), 25 ng/mL FGF7 (Novopro Labs, 318990), 100 ng/mL FGF10. Cultures were fed with fresh expansion medium every 2-3 days, and once organoids grew large enough (∼8-10 days), they were passaged in a 1:3-1:5 ratio. First, Matrigel domes were lifted from the cell culture plates with ice-cold DPBS (Fisher, 10769033), transferred to a 15 mL tube, then mixed thoroughly with a pipette (∼50 times) before being centrifuged (300 x g for 5 minutes at 4°C). After removing the supernatant, 2 mL of TrypLE express was added and the cultures were digested until they dissociated into small clumps (∼4 minutes). Then the cell suspension was mixed thoroughly with a pipette again (∼50 times), then the digestion was stopped by addition of 6 mL of DPBS and centrifuged (300G for 5 minutes at 4°C). The cell pellet was then encapsulated in 50 µL/well of growth factor reduced Matrigel (Corning, 734-1101) and seeded in droplets within a pre-warmed 24-well plate. Cultures were incubated for at least 15 minutes before adding fresh expansion medium containing 10 µM of Y-27632 ROCK inhibitor.

For hormone treatment experiments, organoid cultures were first grown in expansion medium for 5 days before 4 days of differentiation media. Differentiation media contained phenol red free DMEM/F12 with L-glutamate and HEPES (Gibco, 11039021) with 5% conditioned Noggin medium kindly obtained from Dr. Vanessa Muncan and Dr. Gijs van der Brink, 5% conditioned R-spondin 1 (Trevigen, 3710-001-01), 1X B27, 1X N-2 supplement, 1.25 mM N-acetyl-cysteine, 25 ng/mL FGF7, 100 ng/mL FGF10, 1 µM PD153035, 1 µM Troglitazone, and 10 µM retinoic acid. Henceforth, medium only containing phenol red free DMEM/F12 with L-glutamate and HEPES is referred to as base medium.

### Transwell barrier culture

Barrier cultures on 12-well Transwell inserts (Stem Cell, 38023) were prepared by a similar protocol to passaging organoids, except with a longer digestion (6 minutes instead of 4) to allow for smaller clumps and single cells to be formed for seeding. Transwell insert surfaces were precoated with 2% Matrigel in PBS (Applichem, A0965.9010) at 37°C for at least an hour. A total of 1.1 million cells in 300 µL of cell suspension (1 million cells/cm^2^) was seeded onto the apical surface, with 700 µL of medium added to the basal side. Additional medium was provided for the following feedings (500 µL for apical and 1000 µL for basal) every 2-3 days until cultures reached confluency (∼7 days). Once confluence was achieved, the apical medium was replaced with differentiation medium for 3 days. Finally, the basolateral medium was replaced with differentiation medium in the bottom and an air-liquid-interface (ALI) was established by removing all medium from the apical side.

### Hormone exposure

Organoid cultures were exposed to hormones only in diluted differentiation media in the following concentrations: 10 nM β-estradiol (E2, MP Biomedical, 2194565), 1 nM testosterone (Sigma, 86500), and 1 µM norepinephrine (Fisher, AAL0808703). Hormones were aliquoted and frozen to avoid repeated freeze-thaw cycles.

### ELISA assay quantification

Concentrations of CXCL8 (Biolegend, 43504), IL6 (Biolegend, 430504), and TNF-α (Biolegend, 430204) were quantified by sandwich ELISA using commercial kits and the manufacturers’ recommended reagents and protocols. (Biolegend 423501, 421601, 423001). Absorbance at 450 nm and 570 nm (background) were measured with a Spectramax M3 spectrophotometer (Molecular Devices), and the standard curve was with a four-parameter logistic (4PL) model.

### LDH cytotoxicity assay quantification

The cytotoxicity of Transwell barrier cultures was assessed by measuring LDH levels in basolateral supernatant via the CyQuant LDH cytotoxicity assay (Invitrogen, C20300). Absorbance was measured at 490 nm and 680 nm and used for background correction. Percent viability was assessed per manufacturer’s protocol via the experimental LDH activity, maximum LDH activity and blank, which was calculated per hormone group.

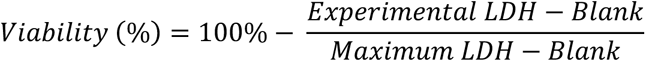

Equation 1 - LDH viability calculated in units of percent, where the difference between the experimental LDH activity and the blank was divided by the difference between the maximum LDH and the blank.

### Immunostaining

Medium was removed, and cultures were fixed with 4% PFA (Fisher, 15434459) at room temperature for 10 minutes, before being washed twice with PBS and permeabilized with 0.1% Triton X-100 in PBS. Next, samples were washed twice with PBS and blocked in blocking buffer containing 2% goat serum (Fisher, NC9660079) and 1% BSA (Fisher, B14) in immunostaining buffer for 1 hour at RT. Immunostaining buffer consisted of PBS with 0.05% Tween20 (Sigma, P1379) and 0.02% sodium azide (Sigma, 26628-22-8). Primary antibodies were diluted in blocking buffer and incubated for 2 h at room temperature or overnight at 4°C (Table 2). Next, cultures were washed four times with immunostaining buffer and when required incubated with secondary antibodies for 1 hour at room temperature or overnight at 4°C. Before imaging, cultures were washed five times with immunostaining buffer and stored at 4°C in 100 µM Trolox (Novus Biologicals, CB-1000-2-NB) in PBS. Fluorescent stains (e.g., Hoechst 33342) were applied together with the secondary antibodies. Imaging was conducted using a Lionheart FX automated fluorescent microscope.

**Table 1.** Hormone concentrations by exposure. Concentrations were informed from previously published clinical data and in vitro experiments.

| Hormone | Concentration |
| --- | --- |
| β-estradiol <sup>6-8,25</sup> | 10 nM |
| Testosterone <sup>6,10,11,26</sup> | 1 nM |
| Norepinephrine <sup>27</sup> | 1 μM |

**Table 2.** Antibody and staining list.

| Target or Product Name | Host | Vendor | Product No | Excitation (nm) | Dilution |
| --- | --- | --- | --- | --- | --- |
| Hoechst 33342 |  | Invitrogen | H3570 | 350 | 1:2000 |
| MKI67 | Rabbit | Invitrogen | PA5-19462 |  | 1:200 |
| Anti-rabbit | Goat | Invitrogen | A32731 | 488 | 1:1000 |

### Sample embedding and immunohistochemistry

Organoid cultures and Transwell barrier cultures were fixed in 4% PFA overnight at 4°C and for 10 minutes at room temperature, respectively. After fixation, samples were stored in PBS at 4°C before being processed for paraffin embedding using a Leica TP1020 automated tissue processor. Before sample processing, organoids were embedded in 1% agarose. Samples were then transferred to 70% ethanol again (1 hour), 80% ethanol (1 hour), 90% ethanol (1 hour), 100% ethanol (30 minutes, repeated 2 additional times), then xylene (VWR, 28973.328) (1 hour, repeated 2 additional times), and finally paraffin (1 hour, repeated once). Following this step, samples were transferred to a container of melted paraffin and degassed in a Shel Lab heated vacuum oven for 3 cycles of 20 minutes, with increasing pressures of 15, 20, and 25 inches Hg. The cassettes containing the samples were transferred into a paraffin bath and embedded with the Medite TBS 88 Tissue Block System. Solidified samples were stored overnight at -20°C before long-term storage at room temperature. Organoid and barrier culture tissue blocks were sectioned using a microtome (Lieca Jung RM2035) at 7 µm and 12 µm thicknesses, respectively. Tissue sections were transferred to a heated water bath (Lauda H2P) then onto glass slides.

Samples for hematoxylin and eosin (H&E) staining were first deparaffinized. In brief, the cultures were treated with Xylene for 3 minutes twice, 100% ethanol twice (3 and 2 minutes), then on 2-minute treatments with decreasing concentrations of ethanol (96%, 90%, 80%, 70%), before two final immersions in demi water for 5 minutes. Next, samples were immersed in Mayer’s Hematein solution (Merck Millipore, 115938) for 7 minutes, then under running tap water for 10 minutes, before staining with eosin. The slides were dipped twice very quickly in 70%, then 96% ethanol, before 3 immersions in 100% ethanol for 2 minutes, and 3 immersions in Xylene for 2 minutes.

### Bulk RNA-seq analysis

RNA was extracted using the RNeasy plus mini kit (Qiagen, 74134) according to the manufacturers protocol. RNA purity and concentration were assessed using Nanodrop and Qubit instruments, respectively, with the Qubit broad range RNA quantification kit used (Thermo, Q10211). Lysate and RNA were stored at -80°C. Library preparation and polyA mRNA sequencing was conducted by Novogene Europe via Illumina platforms.

Raw sequencing data were analyzed using the nf-core/rnaseq pipeline (3.21). The length-scaled gene counts were then analyzed via the DESeq2 package in R.^28^ Fold-change shrinkage was conducted via apeglm (Figure S1).^29^ Genes were considered to be differentially expressed when they had greater than a 0.58 log_2_ fold-change and FDR-adjusted p-value of less than < 0.05. Clusterprofiler was used to conduct gene set enrichment analyses (10^4^ permutations).^30^ Sets between 10 and 200 component genes were included, with pathways considered significant based on an FDR p-value less than 0.05. The enrichplot, org.Hs.eg.db, and DOSE packages were used to conduct supplementary gene pathway analyses.

### Statistics and software packages

All statistical analyses not directly part of the RNAseq analysis were conducted with R (version 4.6.0) using RStudio (Version 2026.04.0+526) and seed 29483 with the packages emmeans, lmtest, MASS, and vegan. dplyr, tidyr, purr, readxl, and stringr were used for data handling. ggpubr, gt, ComplexHeatmap, ggprism, ggplot2, ggsignif, ggfortify, ggsci, ggrepel, patchwork, scales and circlize were used for data visualization. All experiments had at least three experimental replicates (N) with as many technical replicates (m) as feasible. Sample size information is included with each relevant figure and in the supplemental information. (\**p* < 0.05, \*\**p* < 0.01, \*\*\**p* < 0.001).

## Results

### Differentiation of iPSC cultures into urothelial lineage cells expandable as organoids

iPSC clone NAS2 (also known as EDi002-A) was differentiated into definitive endoderm, then toward a urothelial lineage over 18 days using a previously described differentiation protocol (Figure 1A).^24^ Over the course of the protocol, cells changed morphologically (Figure 1C), with increasing cell-cell junctions and multiple layered appearance by the end of the differentiation protocol. Following completion of the initial differentiation protocol, the cell populations were dissociated and encapsulated in Matrigel, where they formed into organoids (Figure 1D-E).

**Figure 1.**
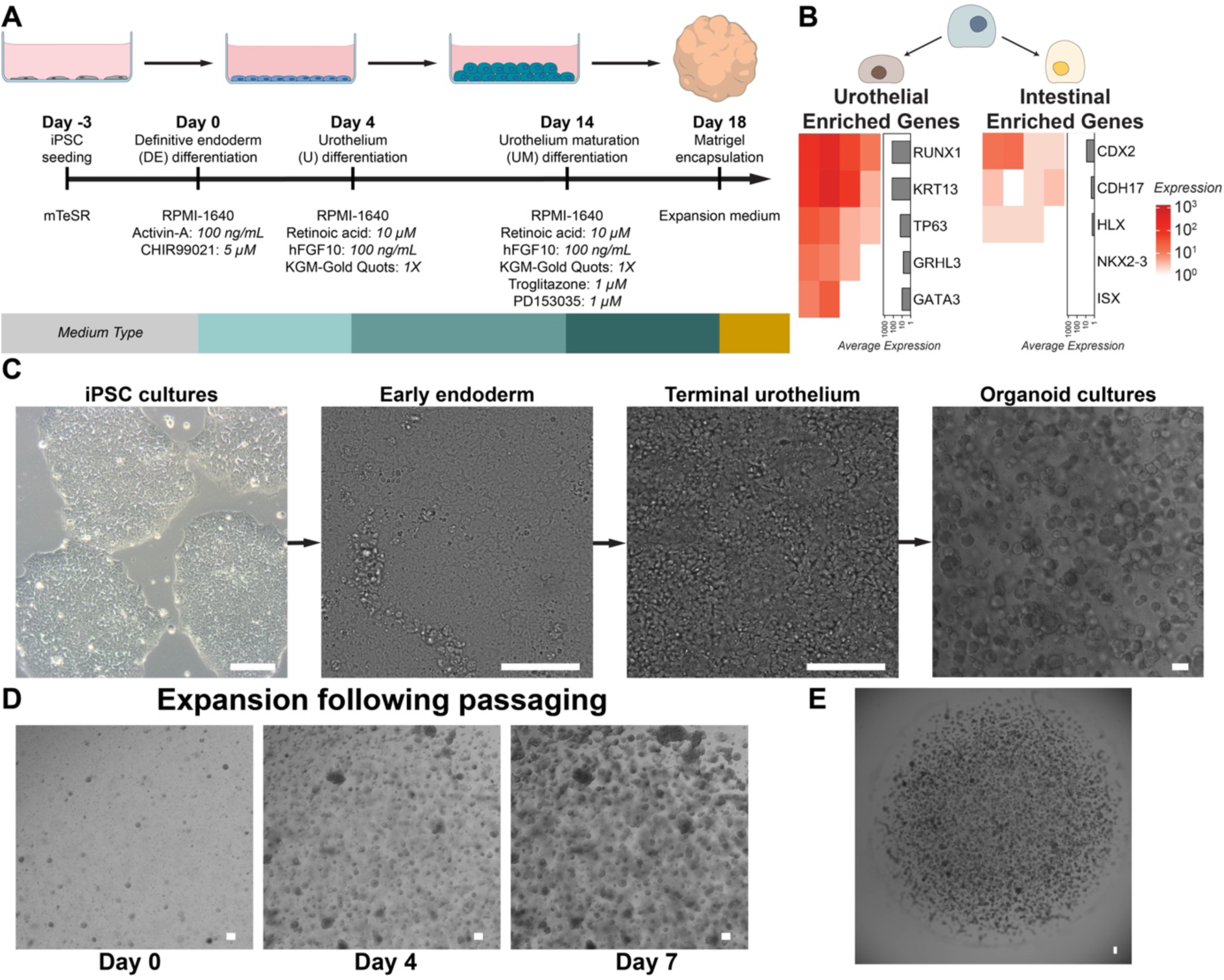
Generation and expansion of iPSC urothelial organoid generation accomplished through three phases of differentiation. A) Schematic representation of iPSC differentiation including key reagents. B) Heatmap of length-scaled transcript counts for genes enriched in urothelial and epithelial lineages in organoids cultured in expansion media. Each column represents a sample. C) Representative images of cells during iPSC differentiation stages. D) Time-lapse images of expansion of organoids following repeated passaging. E) Representative brightfield image of an entire Matrigel dome containing organoids. Scale = 100 µm.

Bulk RNA-seq was performed to characterize gene expression in the resulting organoids. Organoids expressed genes enriched in urothelial lineage cells at much higher levels than those for the developmentally linked, endodermic intestinal epithelium (Figure 1B). Several relevant proteins to urothelial function (Figure 2A) were expressed in organoids, including for transcriptional factors and aquaporins. Genes for key mucosal barrier functions including tight junction components and innate immune response were expressed strongly. The exception was for lipopolysaccharide binding protein (LBP) and TLR7. Several relevant uroplakins and cytokeratins were expressed, however some of the expected types (e.g., UPK1A, UPK2, KRT5, KRT20) were expressed at minimal or nondetectable levels.

**Figure 2.**
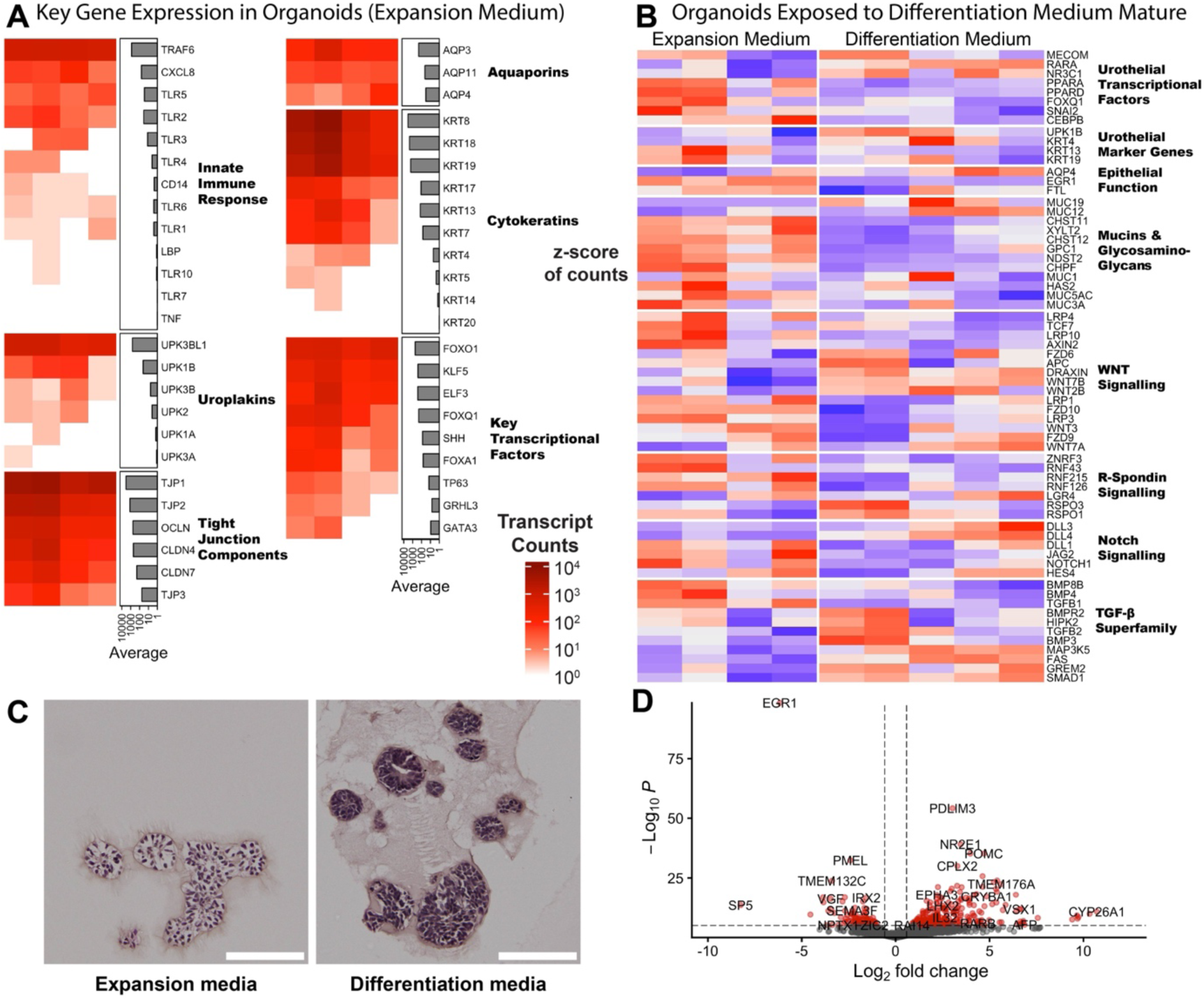
Organoids express several expected genes for functional urothelial cells and are transcriptionally impacted by differentiation medium. A) Heatmap of urothelial relevant length-scaled gene transcript counts in organoids cultured in expansion media, which illustrates the presence of expected genes. Each column represents a sample. B) Select differentially expressed genes (DEGs) from transition of organoids to differentiation media, sorted by their role in urothelial development/function or stem cell maintenance/differentiation. VST transformed counts were z-scaled per gene to visualize variance. C) Representative images of H&E staining of paraffin-embedded cultures with and without introduction of differentiation media. Scale = 100 µm. D) Volcano plot illustrating the most strongly impacted genes from transition of organoids to differentiation media (2859 DEGs). Dashed lines indicate thresholds used for differential gene expression (vertical line describes absolute log_2_ fold-change > 0.58 and horizontal line describes adjusted p-value < 0.05).

Organoids were cultured in differentiation media to promote maturation, which removed select stem cell supporting factors including WNT3A and added several compounds including Troglitazone (PPAR-γ agonist) and PD153035 (EGFR inhibitor), respectively. In addition, retinoic acid was added to stimulate retinoid signaling. Organoids displayed a darker histological appearance following H&E staining (Figure 2C). Bulk RNA-seq analysis identified 2859 differentially expressed genes (DEGs) with a cutoff of an absolute log_2_ fold-change greater than 0.58 and an adjusted p-value less than 0.05 (Figure 2D). Although 50.12% of DEGs were upregulated, among the most confidently impacted genes (adjusted p-value < 10^-6^), 74.69% were upregulated.

Several DEGs relevant to urothelial fate/function and stem cell maintenance/maturation were identified (Figure 2B). Transcription factors associated with retinoid signaling and epithelial/urothelial biology including RXRG, RARA, RARB, PPARA, PPARD, EGR1, and CEBPB were differentially expressed. The uroplakin, UPK1B (log_2_ fold-change = 1.30), was upregulated. The urothelial associated KRT4 (log_2_ fold-change = 1.63) was upregulated, while the general urothelial-associated cytokeratin genes KRT13 (log_2_ fold-change = -2.98) and KRT19 (log_2_ fold-change = -0.95) were downregulated. The aquaporin AQP4 (log_2_ fold-change = 2.07), which is involved in water transport and osmotic regulation was upregulated. Several mucins and glycosaminoglycans (GAGs) were differentially expressed. MUC12 (log_2_ fold-change = 1.82) and MUC19 (log_2_ fold-change = 7.65) were strongly upregulated, while MUC1 (log_2_ fold-change = -0.83) and MUC3A (log_2_ fold-change = -1.83) were downregulated. MUC5AC (log_2_ fold-change = -0.58) was at our fold-change threshold to be considered differentially expressed.

Several genes associated WNT signaling were differentially expressed, including: FZD6 (log_2_ fold-change = 0.66), FZD9 (log_2_ fold-change = -1.01), WNT2B (log_2_ fold-change = 1.33), and WNT7A (log_2_ fold-change = 1.98). Similarly, several DEGs relating to R-spondin signaling were observed, such as: RSPO1 (log_2_ fold-change = 3.23) and LGR4 (log_2_ fold-change = 0.65). Notch-associated genes including DLL4 (log_2_ fold-change = 3.25), and DLL1 (log_2_ fold-change = -0.62). NOTCH1 (log_2_ fold-change = -0.67) just missed the p-value threshold (adjusted *p* = 0.052). Finally, the TGF-β superfamily was also strongly impacted. BMP3 (log_2_ fold-change = 2.34), BMP4 (log_2_ fold-change = -1.64), BMP8B (log_2_ fold-change = -1.11), and the receptor BMPR2 (log_2_ fold-change = 0.61) were differentially expressed. TGFB1 (log_2_ fold-change = - 2.18) was downregulated, but TGFB2 (log_2_ fold-change = 1.24) was upregulated. One of the DEGs with the strongest fold-change expression is NR2E1 showed one of the largest positive fold changes (log_2_ fold-change = 3.46), which has been associated with cellular senescence.

Gene set enrichment analysis using the gene ontology (GO) database identified 19 significantly enriched GO terms (adjusted p-value < 0.05). Ten of the enriched terms were related to retinoid signaling or metabolism were significantly altered (Figure 3A). Shared leading edge genes between three of these gene sets include CRABP2 (retinoic acid binding protein), RBP1 (retinol acid binding protein), and CYP26A1 (retinoic acid degradation enzyme) (Figure 3B, E). Additional enriched terms were associated with chemokine activity/regulation, glycosyl-related hydrolase activity, and neuropeptide/neurotransmitter signaling.

**Figure 3.**
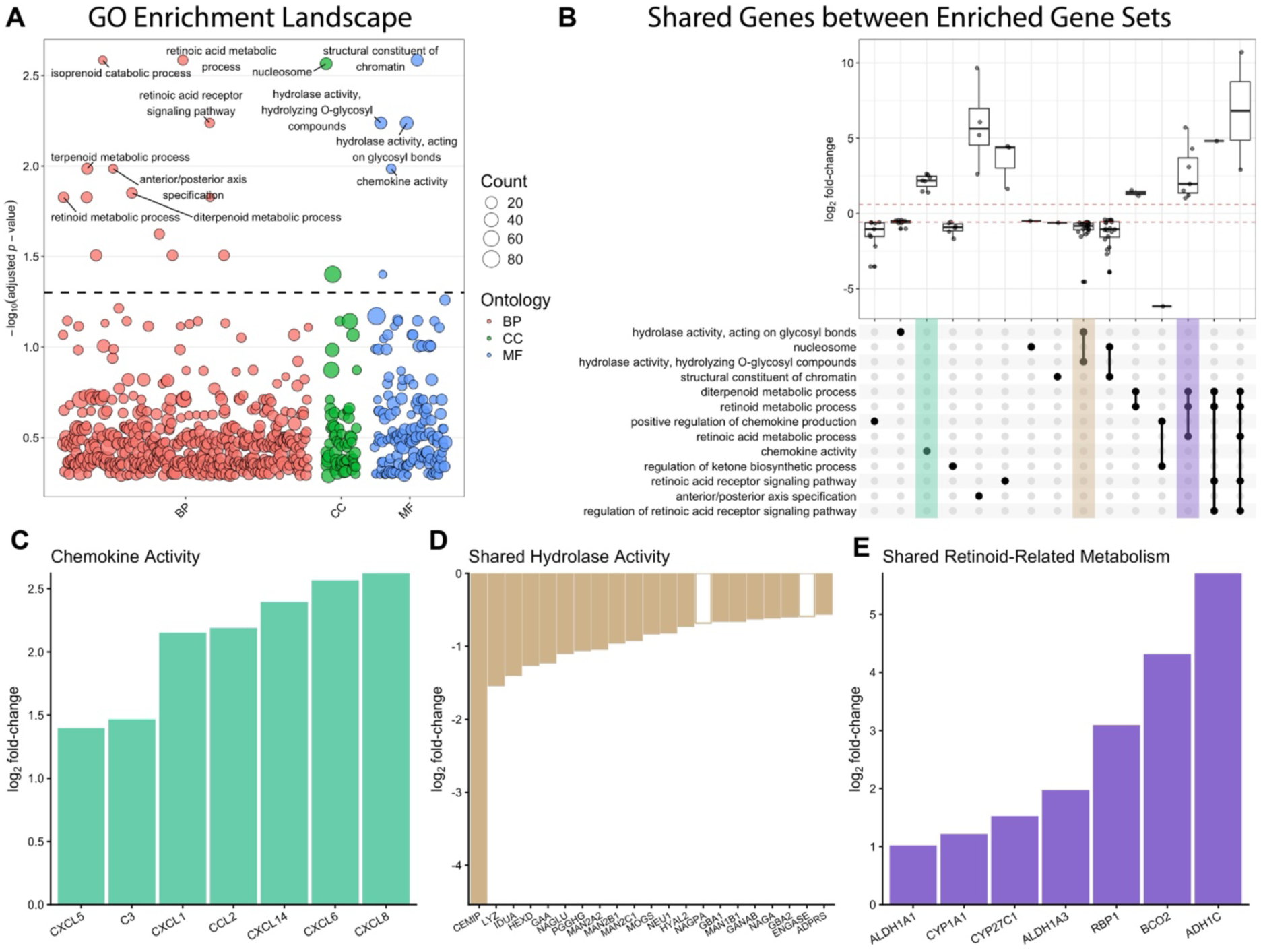
Gene set enrichment analysis of transcriptional changes induced by differentiation medium in urothelial organoids using Gene Ontology (GO) gene sets. A) Manhattan plot of all GO terms separated by ontology type (BP = biological process, CC = cellular component, MF = molecular function). The horizontal dashed line indicates an adjusted p-value of 0.05. B) UpSet plot of the 13 selected significantly enriched GO terms. Boxplots above show the distribution of log2 fold changes for genes represented by each intersection. Barplots show log_2_ fold-change of leading-edge genes from selected gene sets highlighted in the UpSet plot: C) chemokine activity, D) common enriched genes between the gene sets for hydrolase activity acting on glycosyl bonds and hydrolyzing O-glycosyl bonds and E) common genes between the diterpenoid, retinoid, and retinoic acid metabolic process gene sets. Dashed red line in UpSet plot indicates the 0.58 log_2_ fold-change cutoff used for designating genes as differentially expressed. Filled bars indicate genes that were significantly differentially expressed compared with expansion-medium organoids (adjusted *p* < 0.05).

To understand how core enrichment genes were shared between enriched gene sets, 13 significantly enriched GO terms were visualized via an UpSet plot, with log_2_ fold-changes shown for leading edge genes unique or shared among the selected gene sets (Figure 3B). The chemokine activity set (NES = 2.0) contained many chemokine ligands (Figure 3C), such as CXCL8 (log_2_ fold-change = 2.62) and CCL2 (log_2_ fold-change = 2.18), and the complement component C3 (log_2_ fold-change = 1.46). Two hydrolase activity gene sets (Figure 3D) for hydrolyzing O-gylcosyl compounds (NES = -2.14) and for acting on gylcosyl bonds (NES = - 2.03) were downregulated. Between them they shared 21 enriched genes, including the GAG-degraders IDUA (log_2_ fold-change = -1.40) and HEXD (log_2_ fold-change = -1.26), and peptidoglycan hydrolase LYZ (log_2_ fold-change = -1.54). Finally, among the selected retinoid-associated gene sets, those representing diterpenoid (NES = 2.08), retinoid (NES = 2.06), and retinoic acid (NES = 2.13) metabolic processes were positively enriched and shared several genes (Figure 3E), including the carotenoid oxidizer BCO2 (log_2_ fold-change = 4.30) and retinol binding protein RBP1 (log_2_ fold-change = 3.08).

### Estradiol supplementation induces transcriptional changes in differentiated urothelial organoids

To investigate how menopause-associated hormonal changes may influence urothelial biology, female iPSC derived urothelial organoids were supplemented with estradiol, norepinephrine, or testosterone. Hormone concentrations were selected based on clinical data and previous in vitro studies.^5,11,27^ Prior to hormone supplementation, cultures were grown in expansion medium for 5 days (Figure 4A), then cultured in differentiation medium for 4 additional days. The differentiation medium was phenol red free to minimize potential estrogenic effects of phenol red^31^; however both the expansion and differentiation media contained N-2 supplement, which contains progesterone (final concentration = 20 nM). Thus, we did not evaluate the individual impact of progesterone on the urothelium. Estradiol treated organoids displayed visible outgrowths in representative brightfield images (Figure 4B).

**Figure 4.**
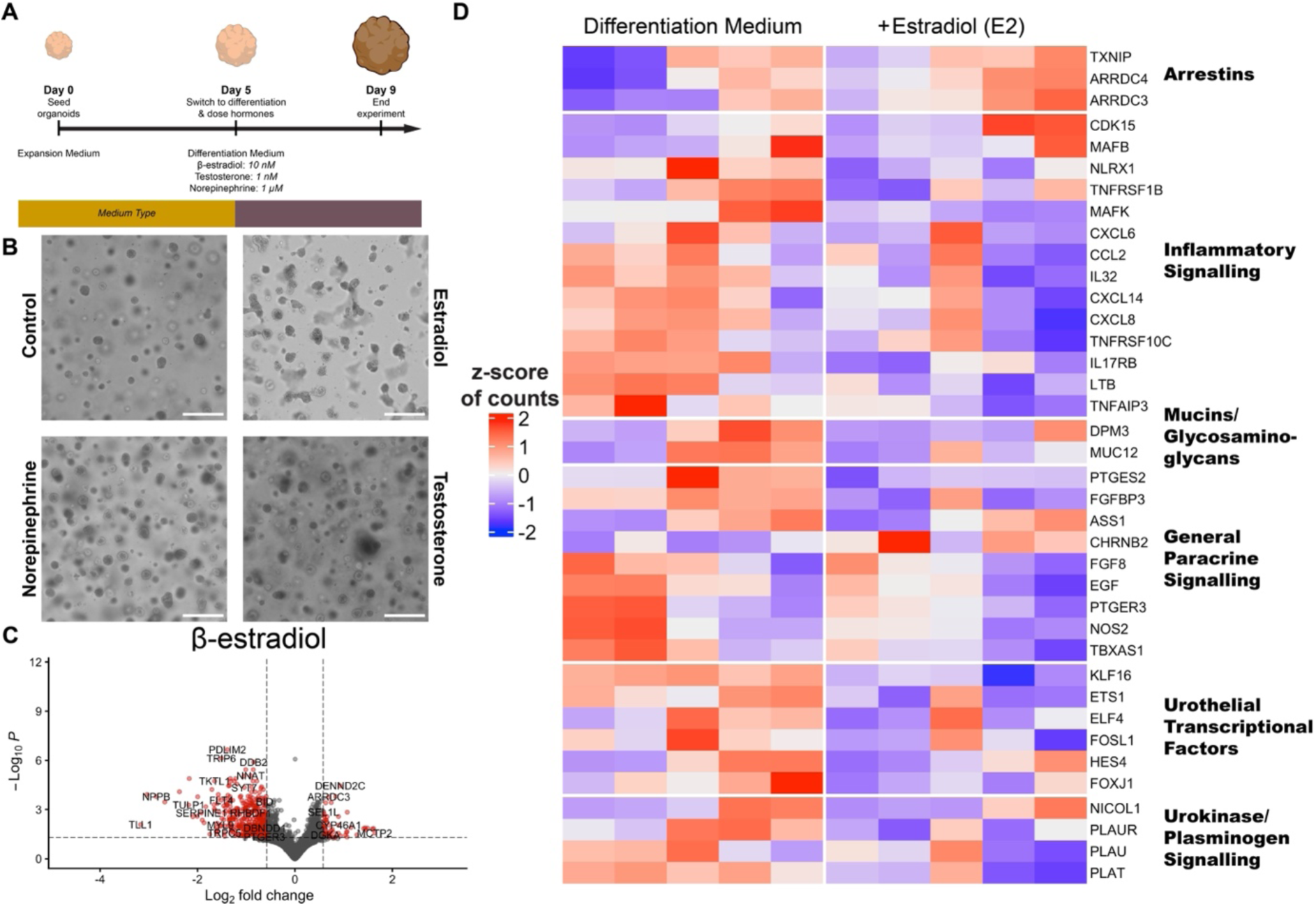
Effects of administration of estradiol, norepinephrine, and testosterone supplementation on gene expression in urothelial organoids. A) Schematic representation of organoid expansion then differentiation followed by differentiation and hormone treatment. B) Representative brightfield images of organoids under each treatment condition (scale = 300 µm). C) Volcano plot of the impact of estradiol treatment on organoid gene expression. Dashed lines illustrate the cutoffs for considering genes differentially expression (vertical line = absolute log_2_ fold-change > 0.58 and horizontal line = adjusted p-value < 0.05). Each dot represents a gene, with red dots indicating a differentially expressed gene (457 DEGs). D) Heatmap of selected genes’ expression organized by general function. Each column represents a sample, with colors determined from the z-score of VST-normalized length-scale gene counts.

RNAseq analysis revealed a markedly different transcriptional response among the hormone treatments. Estradiol supplementation resulted in 457 differentially expressed genes (DEGs) whereas only one DEG was detected following supplementation with norepinephrine (ENSG00000259522), or testosterone (LINC02802) (Figure S1). Applying less stringent thresholds (absolute log_2_ fold-change > 0.38 and adjusted *p* < 0.1) still identified relatively few DEGs (8 for testosterone and 1 for norepinephrine). For this reason, additional sequencing analysis was not completed for testosterone or norepinephrine.

Following estradiol supplementation, the majority of DEGs were downregulated (86.7%, ^F^igure ^4^E). Upregulated genes included the arrestin-domain-containing genes ARRDC3 (log_2_ fold-change = 0.69) and ARRDC4 (log_2_ fold-change = 0.63). In addition, the alpha-arrestin TXNIP (log_2_ fold-change = 0.38) was significantly increased but missed the fold-change threshold. Several genes associated with inflammatory signaling were downregulated (Figure ^4^D), including TNFRSF1B (log_2_ fold-change = -1.20), also known as TNFR2, CXCL8 (log_2_ fold-change = -0.63), and NLRX1 (log_2_ fold-change = -0.76). Additionally, the epithelial mucin MUC12 (log_2_ fold-change = -1.05) and key mannosylation-associated gene DPM3 (log_2_ fold-change = -0.58) were downregulated. Several differentially expressed genes associated with cellular and paracrine signaling were also identified, including EGF (log_2_ fold-change = -0.81), PTGER2 (log_2_ fold-change = -0.62), ASS1 (log_2_ fold-change = -0.59), and NOS2 (log_2_ fold-change = -1.02). Several transcriptional factors were also downregulated as well including FOXJ1 (log_2_ fold-change = -0.62) and ETS1 (log_2_ fold-change = -1.33). Many elements of plasminogen and urokinase signaling were differentially expressed, including PLAU (log_2_ fold-change = -0.60), PLAT (log_2_ fold-change = -0.66), and SERPINE1 (log_2_ fold-change = -1.91). PLAUR was significantly differentially expressed but did not meet the fold-change threshold (log_2_ fold-change = -0.52).

GO gene set enrichment analysis (Figure 5A) resulted in 56 significantly regulated sets (adjusted p-value < 0.05). The top 13 significantly enriched GO terms with the highest absolute normalized enrichment scores (NES) showed relatively limited overlap in their leading-edge genes, although overlap was observed among terms associated with antimicrobial host defence (Figure 5B). Further, the serine-type endopeptidase inhibitor activity gene set was negatively enriched (NES = -1.98). The leading-edge genes (Figure 5C) included several members of the superfamily of serine protease inhibitors (serpins family) such as SERPINE1, SERPINF1 (log_2_ fold-change = -0.58), and SERPINB6 (log_2_ fold-change = -0.52). The protein deglycosylation gene set (NES = 2.05) was positively enriched and included several leading-edge genes (Figure 5D) related to glycoprotein processing and mannose trimming, (Figure 5D) including EDEM2 (log_2_ fold-change = 0.27), EDEM3 (log_2_ fold-change = 0.47), and MAN1A1 (log_2_ fold-change = 0.26). Leading-edge genes also included three genes associated with ubiquitination-mediated protein degradation including RNF103 (log_2_ fold-change = 0.47), RNF139 (log_2_ fold-change = 0.36), and RNF185 (log_2_ fold-change = 0.25). Additionally, the protein demannosylation gene set (Figure S2G) was significantly positively enriched (NES = 1.94), with all leading-edge genes shared with the protein deglycosylation gene set. Estradiol supplementation was associated with negative enrichment of the antimicrobial humoral immune response mediated by antimicrobial peptide gene set (NES = -2.09). Several leading-edge genes associated with term were downregulated (Figure 5E) including CXCL14 (log_2_ fold-change = -1.06) and CXCL6 (log_2_ fold-change = -0.75). To examine AMP gene expression more broadly, genes corresponding to antimicrobial peptides listed in the APD6 Antimicrobial Peptide Database^32^ were assessed following estradiol supplementation. Of these, seven were DEGs (NPPB, POMC, CXCL14, RARRES2, H2AJ, CXCL6, ADM) and all were downregulated (Table S2). Additionally, the enzyme FAP (log_2_ fold-change = -0.99), which processes the AMP NPPB, was also differentially expressed.

**Figure 5.**
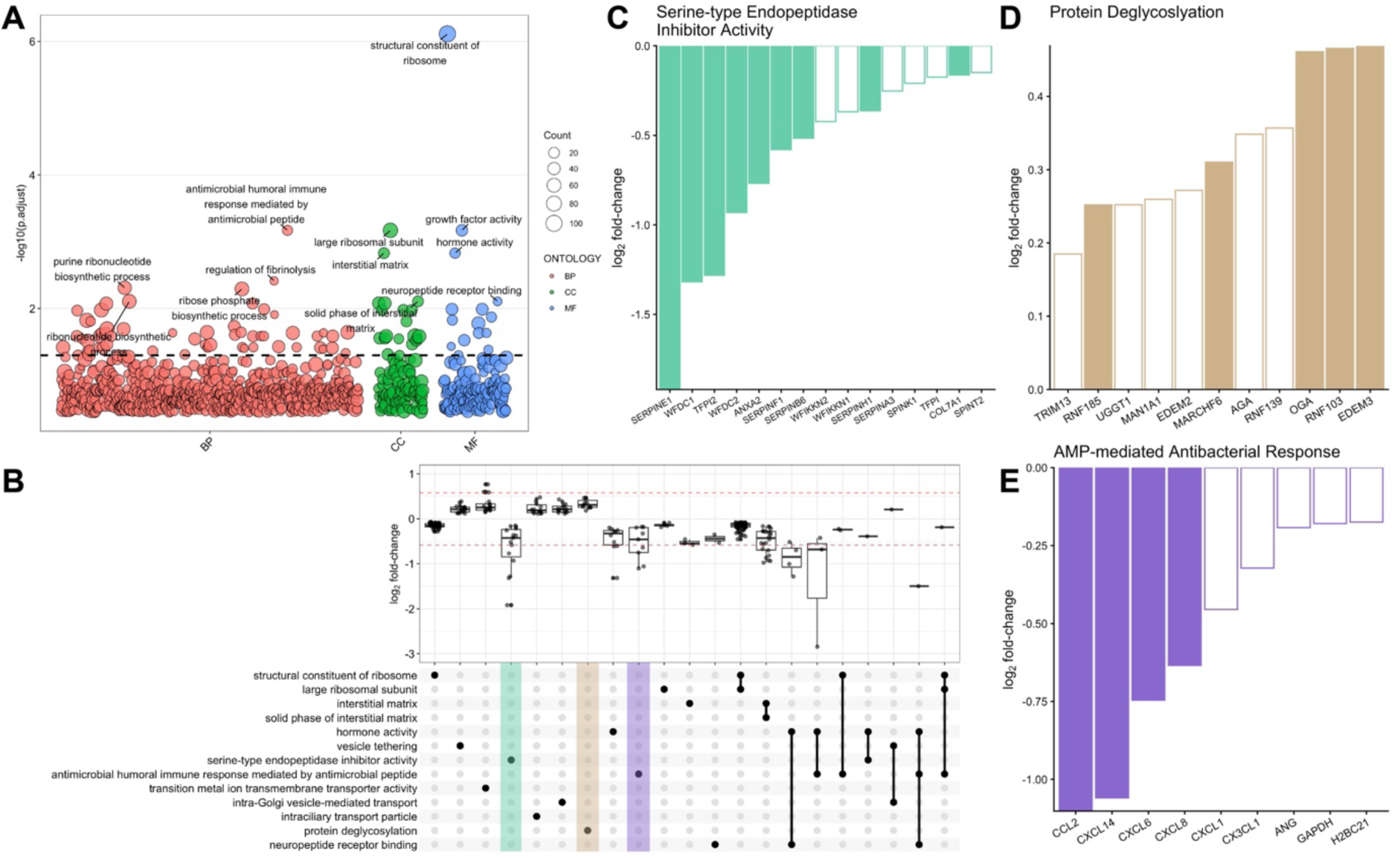
GO gene set enrichment analysis of the impact of estradiol on differentiated urothelial organoids. A) Manhattan plot of all GO terms plotted against log-scaled adjusted p-value. Dotted black line indicates the 0.05 threshold. B) UpSet plot of shared leading-edge genes from the top thirteen gene pathways including boxplots of the log_2_ fold-change expression. Horizontal dashed red lines indicate the log_2_ fold-change cutoff of +/-0.58 for differential expression. Barplots of the log_2_ fold-change expression of leading-edge core enrichment genes within pathways highlighted in the enrichment map and UpSet plot: C) Serine-type protease inhibitor activity, D) Protein deglycosylation, E) Antimicrobial humoral immune response mediated by antimicrobial peptide (AMP). Bars without a fill color were not significantly expressed compared to differentiation medium-only controls (adjusted *p* < 0.05).

The growth factor activity gene set was negatively enriched (NES = -1.98), with 23 leading edge genes (Figure S2B). These included EGF, but also VEGFB (log_2_ fold-change = -0.75), VEGFC (log_2_ fold-change = -0.56), and the NOTCH ligand JAG2 (log_2_ fold-change = -0.58). The transition metal transmembrane transporter activity gene set was positively enriched (NES = 1.98), with 16 leading-edge genes including the iron-exporting transporter SLC40A1 (log_2_ fold-change = 0.59), zinc antiporter SLC30A4 (log_2_ fold-change = 0.60), and copper SLC46A3 log_2_ fold-change = 0.77). Mouse Slc46a3 has been shown to enhance cellular responses to peptidoglycan-derived NOD agonists, suggesting that human SLC63A may have a similar function.^33^ The gene set for regulation of fibrinolysis (NES = -1.96) was negatively enriched, with contributions from the previously mentioned plasminogen and urokinase signaling genes (Figure S2F).

### Evaluation of the maturation of urothelial barrier cultures on Transwells following differentiation and introduction of air-liquid interface

We aimed to finally understand how urothelial barrier culture expression on a Transwell compared to organoids. To do this, organoids were first dissociated and seeded onto a semi-permeable Transwell membrane, whereby the cultures received expansion medium for seven days to allow for a confluent layer to form (Figure 6A). Next, for three days, a differentiation gradient was established, where the medium in the apical compartment was replaced with differentiation medium, working from the principle that differentiating the apical cell layers while maintaining the proliferative signals to the basolateral-most layers better mimics native tissue cues. Finally, air-liquid interface (ALI) was established by removing all medium from the apical compartment for 3 days, with the basolateral side compartment medium being replaced with differentiation medium. Transwell cultures formed a confluent and multilayer barrier with MKI67-positive cells (Figure S5).

**Figure 6.**
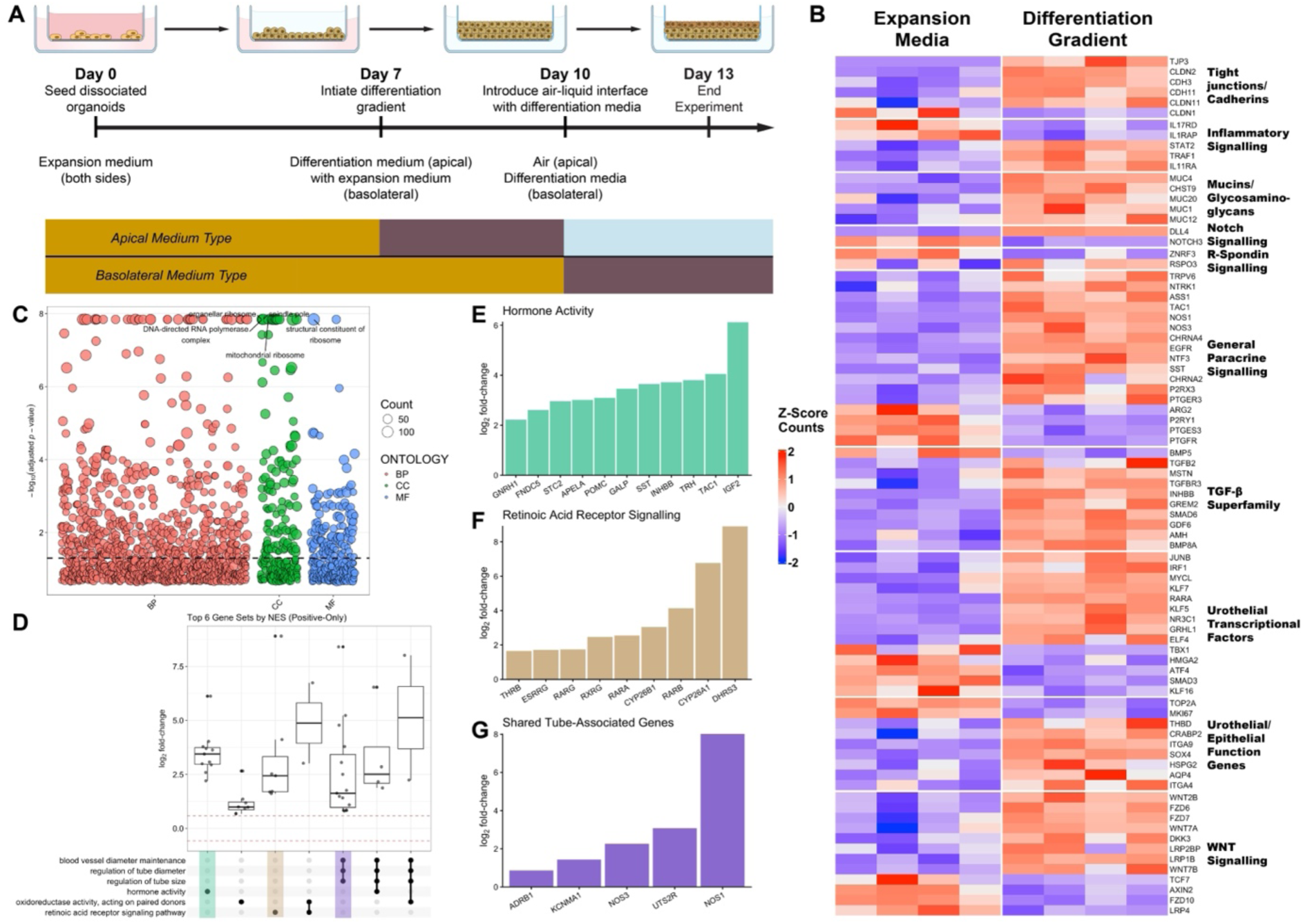
A) Barrier cultures established from dissociated urothelial organoid cultures were cultured for 7 days in expansion medium then transitioned to a differentiation gradient and air-liquid interface culture. B) Heatmap of selected genes populated from the z-score of VST-transformed count data. C) Manhattan plot of GO terms from gene set enrichment analysis of the impact of differentiation gradient on expansion cultures. Black dotted line annotated at an adjusted p-value of 0.05. D) UpSet plot of shared enriched genes between the top six positive normalized enrichment score gene sets. Log_2_ fold-change gene expression of core composed genes per comparison included above with red dotted lines indicating the fold-change cutoff for differential expression (log_2_ 0.58). Among the top six GO terms by positive NES, enriched genes from the E) hormone activity and F) retinoic acid receptor signaling gene sets alone, and G) shared between the blood vessel diameter, regulation of tube diameter, and regulation of tube size gene sets. Solid filled bars indicate genes which were below an adjusted p-value of 0.05.

Bulk RNA-seq analysis of these cultures resulted in 3317 DEGs (Figure S3A). Of these, 71.7% were upregulated. Upregulated genes included a very high number of tight junctions and cadherins (34 DEGs, Figure 6B), such as TJP3 (log_2_ fold-change = 6.01), CDH3 (log_2_ fold-change = 1.08) and CDH11 (log_2_ fold-change = 1.72). Several inflammatory signaling genes were also impacted including IL1RAP (log_2_ fold-change = -1.00), STAT2 (log_2_ fold-change = 0.61), and TRAF1 (log_2_ fold-change = 1.17). MUC1 (log_2_ fold-change = 1.49), a major component of the urothelial mucin environment was upregulated, along with MUC4 (log_2_ fold-change = 3.37) and MUC12 (log_2_ fold-change = 1.36). Few Notch and R-Spondin genes were differentially expressed. NOTCH3 (log_2_ fold-change = -0.76) was one exception. RSPO1 (log_2_ fold-change = 0.66) was slightly above the significance threshold (adjusted *p* = 0.08). There were a large amount of upregulated genes related to various paracrine signaling systems: nitric oxide via NOS1 (log_2_ fold-change = 8.01) and NOS3 (log_2_ fold-change = 2.24), ATP via P2RX3 (log_2_ fold-change = 1.50), serotonin via HTR1A (log_2_ fold-change = 8.41) and HTR1B (log_2_ fold-change = 5.23), arginine via ASS1 (log_2_ fold-change = 2.69) and ARG2 (log_2_ fold-change = -0.62), and prostaglandins via PTGES3 (log_2_ fold-change = -0.59) and PTGER3 (log_2_ fold-change = 1.46). Several genes from the TGF-β superfamily, including 12 BMP genes, were differentially expressed including TGFB2 (log_2_ fold-change = 3.07), TGFBR3 (log_2_ fold-change = 0.74), and BMP8A (log_2_ fold-change = 1.05).

Among urothelial-associated transcriptional factors, many were also differentially expressed including JUNB (log_2_ fold-change = 0.63), TBX1 (log_2_ fold-change = -0.90), and ATF4 (log_2_ fold-change = -0.81). Specific to urothelial function and cell type markers, MKI67 (log_2_ fold-change = -0.74), TOP2A (log_2_ fold-change = -0.71), and AQP4 (log_2_ fold-change = 2.51). Finally, several WNT-signaling associated genes were differentially expressed including FZD6 (log_2_ fold-change = 0.91), FZD7 (log_2_ fold-change = 0.62), WNT2B (log_2_ fold-change = 0.93), WNT7A (log_2_ fold-change = 1.04), and WNT7B (log_2_ fold-change = 1.17).

To understand how gene sets may be regulated, gene set enrichment analysis using GO terms was again conducted. 800 gene sets were significantly enriched (Figure 6C). The gene sets with the largest absolute NES were dominated by downregulated gene sets (72.8%), which often related to ribosomal pathways (Figure S3C). To extract more unique information from this analysis, we also explored the top upregulated, or positive NES, gene sets (Figure 6D). The hormone activity gene set (NES = 2.01) resulted in 11 genes unique to the other sets in the UpSet plot (Figure 6E) such as the substance P producer TAC1 (log_2_ fold-change = 4.03) and the somatostatin hormone producer SST (log_2_ fold-change = 3.63). Additionally, the retinoic acid receptor signaling pathway was upregulated (NES = 1.98) with several retinoic acid receptors (Figure 6F) including RARB (log_2_ fold-change = 4.11) and RXRG (log_2_ fold-change = 2.44). Three upregulated gene sets (Figure 6G) relating to tube morphology including regulation of tube size (NES = 1.98) and diameter (NES = 1.98) and blood vessel diameter maintenance (NES = 1.98) shared 15 enriched genes. For example, the urotensin II receptor UTS2R (log_2_ fold-change = 3.63).

In addition, the nitric oxide mediated signal transduction (NES = 1.97) and metabolic process (NES = 1.67) gene sets were also significantly upregulated (Figure S3E-F), with the unique enriched genes including THBS1 (log_2_ fold-change = 2.74) and EDN1 (log_2_ fold-change = 2.16). NOS1 and NOS3 were the only shared enriched genes between the sets. Additionally, twenty-five genes were enriched in the neurotransmitter receptor activity set (NES = 1.91, Figure S3G), with many acetylcholine receptor subunits for example CHRNA2 (log_2_ fold-change = 3.69) and CHRNA4 (log_2_ fold-change = 2.38).

After three days of ALI culture, the urothelial barrier cultures had 1088 genes differentially expressed (Figure 7A, Figure S3B), with 33.9% upregulated. Among these were the tight junction or cadherin genes DSG2 (log_2_ fold-change = -0.88) and JAM2 (log_2_ fold-change = 0.62). In addition, the inflammatory signaling genes including the flagellin-detecting TLR5 (log_2_ fold-change = -1.82) and NOD-like receptor NLRX1 (log_2_ fold-change = -1.13). Mucins and GAG-related genes such as MUC4 (log_2_ fold-change = -1.38). Notch and TGF-β superfamily genes were differentially expressed, excluding, for example, DLL3 (log_2_ fold-change = 1.30) and SMAD7 (log_2_ fold-change = 1.08), respectively. The paracrine signaling genes for the inflammatory neuropeptides (e.g., substance P) were differentially expressed including TAC1 (log_2_ fold-change = 0.77), TACR1 (log_2_ fold-change = 1.37), and TACR3 (log_2_ fold-change = - 0.65). NOS1 (log_2_ fold-change = 1.45) and ASS1 (log_2_ fold-change = 2.51) were upregulated. The urothelial-associated transcriptional factors JUN (log_2_ fold-change = 1.06) and NR2F6 (log_2_ fold-change = 0.86). The urothelial proliferative cell marker MKI67 (log_2_ fold-change = 0.56, adjusted *p* = 2.5 x 10^-5^) missed the fold-change cutoff. Several WNT signaling genes were differentially expressed, including LRP2 (log_2_ fold-change = 0.71) and its binding protein LRP2BP (log_2_ fold-change = -0.73). Additionally, the iron transporter SLC11A2 (log_2_ fold-change = 0.32) missed the fold-change threshold.

**Figure 7.**
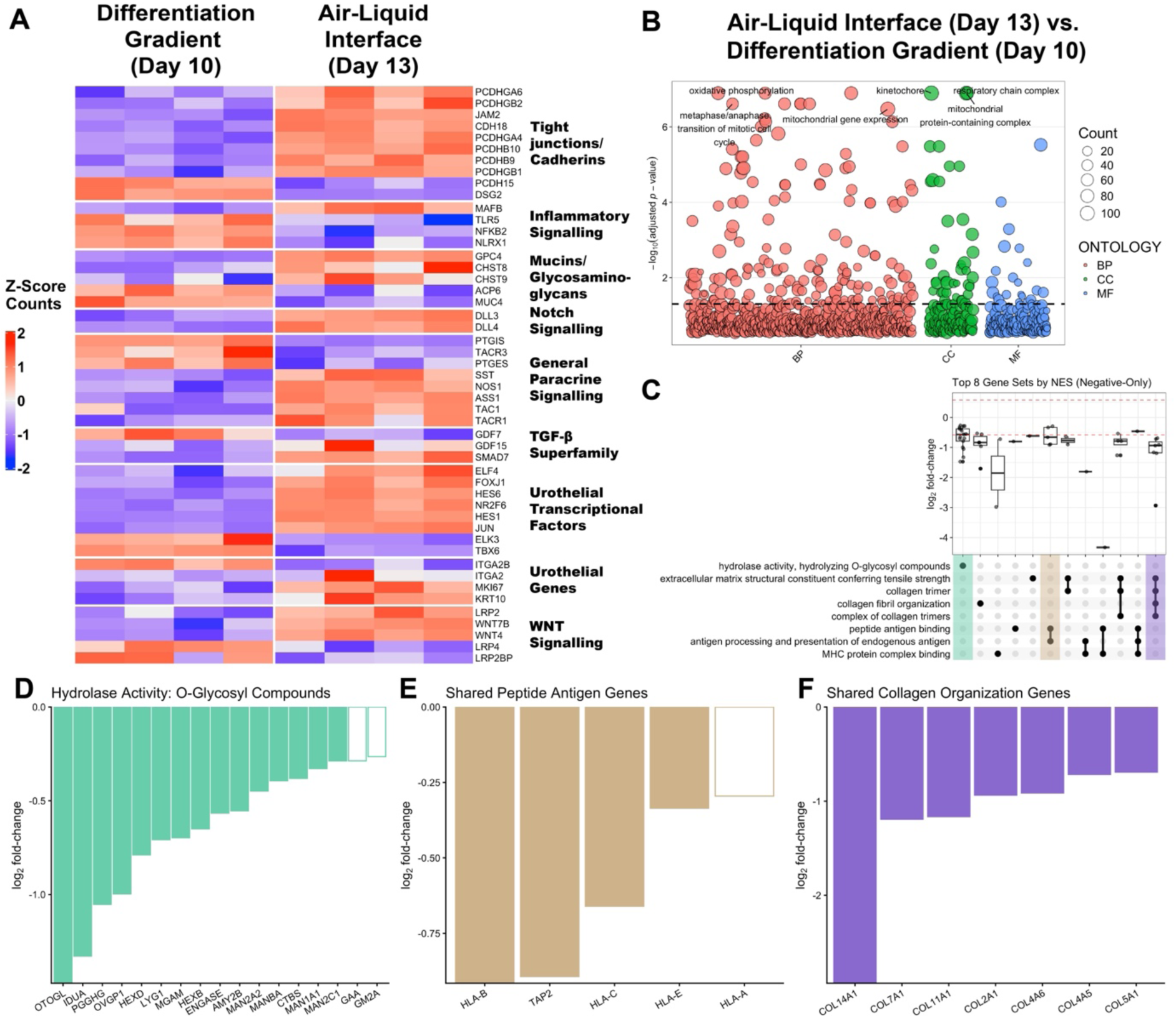
Impact of the introduction of an air-liquid interface on urothelial barrier cultures. A) Heatmap of select genes, with the z-score of the VST-transformed transcript counts. B) Manhattan plot of GO terms from gene set enrichment analysis. C) UpSet plot of top eight negative normalized enrichment score gene sets. Boxplots of the log_2_ fold-change expression provided per unique combination. Dotted red line indicates the 0.58 threshold used to classify genes as differentially expressed. Barplots of the log_2_ fold-change expression of leading-edge core enrichment genes within gene sets: D) Hydrolase activity: O-glycosyl compounds alone, E) shared between peptide antigen binding and antigen processing and presentation of endogenous antigen, and F) shared between extracellular matrix structural constituent conferring tensile strength, collagen trimer, collagen fibril organization, and complex of collagen trimers. Solid filled bars indicate genes which were below an adjusted p-value of 0.05.

Following gene set enrichment analysis, 248 GO terms were significantly enriched (Figure 7B), with 89.9% upregulated (positive NES). The positive NES genes sets were dominated by upregulated cellular respiration functions composed of leading-edge genes with differential expression predominately below the 0.58 fold-change threshold (Figure S4D-E). Again, to identify additional relationships the top gene sets were split directionally (Figure 7C). The hydrolase activity, O-glycosyl compounds gene set was downregulated (Figure 7, NES = - 1.90) with genes including IDUA (log_2_ fold-change = -1.33) and HEXD (log_2_ fold-change = - 0.79). 5 genes were shared between the peptide antigen binding (NES = -1.97) and the antigen processing and presentation of endogenous antigen (NES = -1.92) gene sets (Figure ^7^E), including multiple HLA subunits such as HLA-B (log_2_ fold-change = -0.91) and HLA-C (log_2_ fold-change = -0.66). 7 genes were shared among four gene sets relating collagen fibril organization (NES = -1.96), collagen trimer (NES = -2.07), complex collagen trimer (NES = - 1.94) and extracellular matrix structural constituent conferring tensile strength (NES = -2.08). Not included but enriched, were the gene sets for SMAD protein signal transduction, relevant for TGF-β signaling (NES = 1.87), cytokine receptor activity (NES = -1.80), and antiviral innate immune response (NES = -1.84).

Across the comparisons completed, the effect of differentiation medium on the cultures was had the most unique DEGs observed (Figure S6A). Of the total 3317 DEGs in the Transwell comparison, 16.5% were shared with the organoid expansion-differentiation DEGs. Another 11.0% were shared with the ALI group DEGs. In order to understand how differences in gene expression tell a wider story about our iPSC urothelial organoid model, we conducted a redundancy analysis on all of the bulk RNA-seq results (Figure S6B). By linearly regressing datapoints along RDA ordination space based on the experimental group, redundancy analysis allows for interpretation of distances between points and statistical testing of parameters. Input data was VST transformed via DESeq2 and regressed against experimental group. Conditioning the data for experimental replicate did not improve the explained variance or model fit, thus only experimental group predictors (culture type, differentiation medium, ALI, estradiol) were included as model terms. The overall model and 2 RDA axes were significant (Table S4), plus the effect of experiment culture type (Transwell vs. Organoid) and differentiation medium. 60.0% of the total variance was explained by the constrained axes (RDA1-4). RDA1 (45.5% variance) predominately separated the difference in culture type and RDA2 (10.2% variance) separated the difference in differentiation medium and, to a lesser degree, experimental groups.

To explore how previously identified enriched gene sets directionally ordinated, seven GO terms were selected based on previous gene set enrichment analyses and annotated in the RDA space (Figure S6B-C). Vectors were determined by taking the average loading of the genes within each set. Antimicrobial humoral immune response mediated by antimicrobial peptide was the strongest pathway in RDA1, while retinoic acid receptor signaling pathway and the nitric oxide mediated signal transduction gene sets were the highest in RDA2. Among the genes with the highest loading in RDA1 were three cytokeratins KRT8, KRT18, and KRT19 (Table S5). In RDA2, the highest loadings included two retinoic acid related genes RBP1 and RARB, and the nitric acid gene NOS1 (Table S6).

## Discussion

### Generation of iPSC urothelial cultures

In this work, we describe a novel methodology to generate urothelial organoids from iPSCs and additionally to culture them as a multilayered barrier culture on Transwells. The initial iPSC differentiation protocol matches previously published methods to generate adherent urothelial cells expressing KRT13, KRT20, and uroplakins^24^, which are then dissociated and expanded as organoids. These organoids express many of the expected genes for a functional urothelial barrier (Figure 2B), including key transcriptional factors such as FOXA2, SHH, ELF3, and GATA3. Additionally, they express basal markers of proliferative stem cells required for maintenance including ITGA6 (CD49f), CD44, TP63, KRT4, and MKI67.^34^ Qu et al. recently generated urothelial organoids from iPSCs via the activation of WNT and BMP signaling in hindgut cultures.^22^ They reported that BMP2 treatment alone resulted in a heterogenous culture balanced between intestinal and urothelial positive cells represented by CDX2 and GATA3, respectively. BMP and WNT co-treatment separated cultures from an intestinal lineage via bulk RNAseq, where it expressed key markers roughly ordinated to bladder tissue. Similarly, we observe minimal expression of genes enriched in intestinal lineage cells compared to those for urothelial cells (Figure 1B).

Cytokeratin genes are often used as a marker of urothelial subtypes including basal, superficial, and stem cells. In our differentiated organoids, urothelium-wide cytokeratins are present (e.g., KRT19, KRT18, KRT17) however, not all of those often used as markers for subtypes are (e.g., KRT20, KRT5). Similarly, while some of the uroplakin genes associated with superficial cells are not consistently expressed (UPK1B), others are poorly expressed (UPK1A). It is unclear whether these results are related to a factor with urothelial differentiation (e.g., maturity) or if it reflects the patient biology tested. It is also possible that in combination with the previous conditions, the sequencing depth was insufficient to capture the low expression of uroplakins within a subset of cell populations. Mullenders et al. generated human and mouse urothelial organoids from fresh tissue, however also reported no expression of the superficial cell markers Upk3A or Krt20 in primary healthy murine bladder organoids.^20^ Jasmine and Baraiya et al. recently demonstrated in developing mice (E12.5-E18.5) that uroplakin expression varies between urinary tract segment (bladder vs. urethra), and that female mice have significantly lower (adjusted *p* < 0.05) expression of Upk3BL, Upk3B, Upk2, Upk1A, and Upk1B in the urethra than male mice.^35^ In addition, well described urothelial genes including Foxa1, Krt4, and Shh were differentially downregulated. With spatial transcriptomics on intact mouse bladder tissue, Matković et al. did not identify strong Krt20 expression in their cluster identified as umbrella cells (∼14%), and did not propose it or any of the major uroplakins (Upk1A, Upk1B, Upk2, Upk3) as umbrella cell markers.^36^ Horsley et al. found exposure to pooled patient urine for 15 days resulted in maturation of reported HBLAK cell line barrier cultures, including expression of uroplakins and KRT20.^15^ This motivates future work to explore whether organoid-derived barrier cultures on Transwells exposed to synthetic urine (simUrine)^37^ would demonstrate altered differentiation conditions.

Several works have expanded this understanding with single-cell omics methods on tissue or reviewing existing datasets have aimed to more substantially characterize the urothelium (both urethra and bladder). Jasmine and Baraiya et al. identified several markers of the developing urethra and bladder with bulk RNA-sequencing, confirmed with in situ hybridization. Markers of the urethra included Bcl2, Arg1, Wnt6, and Lgr5. For the bladder, Krt7, Sprr1A, Pparg, Aqp3, Lgals3 were proposed as markers.^35^ Additionally, they identified Irx1, Irx2, Sox2, and Pax9 as urethra-specific transcriptional factors. In our organoids cultured in expansion medium, we observe more genes expressed relating to urethra tissue than the bladder (Figure S7A). They also reported that in the urethra specifically, there were many genes differentially expressed in female vs male mice (1733 with different cutoffs), including Esr2 (estrogen receptor), Fgf3, and Mmp7. Among the DEGs they observed with greater than an absolute fold-change of 16 times (absolute log_2_ fold-change > 4), we observe stronger expression of human analog genes associated with the female urethra (Figure S7B-C). This supports the relevance of studying UTIs, which predominately impact women^2^, with iPSC lines from female donors. Previous work has demonstrated that expressed murine urothelial genes vary from human, so further work investigating human tissue is warranted.^38^

Ramal et al. reviewed and re-analyzed public RNA-seq and ATAC-seq datasets to identify transcriptional factors relevant to cell identities within the human bladder.^39^ They stratified them on whether they were previously established, suggested, or novel. Within those identified, we observe expression of these transcriptional factors in our organoids (Figure S8A), with the weakest expressed being luminally associated. Specifically, Transwell barrier culture differentiation demonstrated a clear pattern of increasing luminal transcriptional factor expression along the length of the experiment (Figure 11). Some of these genes were also differentially expressed following treatments. For example, the luminal-associated transcriptional factors MYCL (log_2_ fold change = 0.65) and NR2F6 (log_2_ fold change = 0.87) were differentially upregulated when a differentiation gradient and ALI were established, respectively. Expression broadly suggests relevant transcriptional factor expression is present, however terminal differentiation genes are not as highly expressed. Nevertheless, establishment of a differentiation gradient in Transwells, then introduction of ALI culture increases luminal transcriptional factor expression. In airway epithelial organoid cultures ALI must be maintained for a period of weeks to properly establish a mature barrier phenotype.^40^ Longer-term ALI and artificial urine culture was tolerated with robust barrier function, thus further exploration of longer-term differentiation in air-liquid interface culture or with artificial urine may yield more mature barrier cultures.

Single-cell RNAseq analyses have proposed additional urothelial cell types including Liu et al. who identified Plxna4-positive superficial cells (Krt20-negative) and Aspm-positive basal cells which they theorize to be involved in wound repair and stem cell proliferation. In our urothelial organoid cultures in expansion medium, we observed expression of both PLXNA4 (average counts = 46.8) and ASPM (average count = 3002.3) consistent with the proposed cell type presence. PLXNA4 does not rise in organoid cultures exposed to differentiation medium (log_2_ fold-change = 0.08), however is differentially upregulated in Transwell cultures in differentiation gradient (log_2_ fold-change = 3.37) and ALI culture (log_2_ fold-change = 0.97). In Transwell cultures, ASPM is differentially downregulated when exposed to the differentiation gradient (log_2_ fold-change = -0.74) and upregulated in ALI culture (log_2_ fold-change = 0.44). In addition, with single-cell RNAseq Jasmine and Samtiya et al. identified seven clusters of cells in the murine urethra, and proposed markers for each type, including urethral neuroendocrine cells (UNECs) and glandular cells.^41^ Although they demonstrated luminal Upk1a expression with in situ hybridization probes, it was not one of the six proposed marker genes from their single-cell RNAseq analysis. All the human analogs of the proposed luminal markers were expressed in organoids (KRT8, KRT18, CLU, MMP7, PIGR), except for CXCL17 (Figure S12). Glandular cells were identified with Kcnma1, Cldn10, and Aqp5. The calcium-activate potassium channel KCNMA1 is differentially upregulated in differentiated organoids (log_2_ fold-change = 1.45) and both differentiated Transwells (log_2_ fold-change = 1.41) and ALI cultured Transwells (log_2_ fold-change = 0.65). No differential expression differences in CLDN10 and minimal AQP5 transcripts were observed in our cultures. Aquaporin and claudin urothelial expression varies dramatically between species, with for example different aquaporin proteins primarily expressed in rat, pig, and human tissue.^38^ UNECs are thought to both sense and communicate with local neural populations.

The urothelium communicates with mesenchymal and neural populations via several molecules including nitric oxide (NO), prostaglandins, purines (e.g., ATP), and acetylcholine.^38^ Such signaling is established as important for orchestrating bladder development^38^, however has been also demonstrated to be involved in innate immune response.^42^ Ambrogi et al. showed UNECs act as sentinel cells and in response to detected LPS secrete serotonin and initiate contraction to flush invading pathogens. Similar to intestinal enteroendocrine cells, UNECs can be identified by ChgA expression, where differentiation media increased expression in organoids (log_2_ fold-change = 1.07) and Transwells (log_2_ fold-change = 0.76). Jasmine and Samtiya et al. also identified Tph1 and Calca as secondary markers. The epithelial serotonin producer TPH1 was expressed on average in all conditions without differential expression, however, was on average higher in Transwells (85 counts) vs. organoids (15 counts). There was minimal CALCA expression present in our cultures. Specified staining and functional testing would be required to identify whether UNECs are also present in our cultures. In the intestine, although enteroendocrine cells compose only around 1% of the epithelial population, they are further specialized by individual signal types, where serotonin-producing enterochromaffin cells are among the most plentiful, with others producing compounds like somatostatin (D-cells) and neurotensin (N-cells) are more rare.^43^

Retrograde neural tracing studies in female pigs have demonstrated 65% of neural projections to the bladder stained positive for somatostatin.^44^ Projections which reached the urothelium were mostly immunoreactive to the acetylcholine transporter SLC18A3. Here, we identify somatostatin signaling and reception to be differentially impacted by culture conditions. The somatostatin production gene SST was increased in both Transwell conditions: the differentiation gradient (log_2_ fold-change = 3.63) and ALI culture (log_2_ fold-change = 2.13). The SSTR2 receptor was differentially expressed in differentiated organoids (log_2_ fold-change = 1.55) and missed the fold-change threshold in differentiated Transwell cultures (log_2_ fold-change = 0.50).

Organoid cultures endogenously express many relevant signaling genes (Figure S10A), including for nociceptive neuropeptides (e.g., TAC1 for isoforms such as Substance P) and neurotrophic factors (e.g., NTRK3 for TrkC). Differentiation of organoids increases expression of urothelial paracrine signaling genes, especially in Transwell cultures, including those associated with nitric oxide, acetylcholine, and purines (Figure S6B, Figure S10B, Figure S11). The neuropeptide signaling pathway is upregulated (Figure 8-9, Figure S6) in differentiated organoids (NES = 2.06, adjusted *p* = 0.018) and Transwells (NES = 1.82, adjusted *p* = 0.006). The differentiated Transwell cultures were upregulated in the nitric oxide mediated signal transduction (NES = 1.96, adjusted *p* < 0.001) and synaptic transmission, cholinergic (NES = 1.78, adjusted *p* = 0.015) gene sets. Nitric oxide, acetylcholine, and purine signaling participates in coordinating bladder function associated with stretching (purines) and relaxation (nitric oxide), however are thought to potentially play larger roles in tissue function.^38^ Transwells exposed to differentiation gradients have large upregulation of NOS1 (log_2_ fold-change = 8.01) and NOS3 (log_2_ fold-change = 2.24). NOS1 is further upregulated with ALI is introduced (log_2_ fold-change = 1.45). Actylcholine changes include CHRNA3 which is upregulated by differentiation media in organoids (log_2_ fold-change = 2.49) and Transwells (log_2_ fold-change = 0.50). Similarly for CHRNB4 in organoids (log_2_ fold-change = 2.04) and Transwell (log_2_ fold-change = 0.56) there was gene upregulation. Estradiol exposure increased CHRMB2 expression (log_2_ fold-change = 0.58). It’s possible that the development of a multilayered barrier in Transwells may have influenced the larger expression differences in signaling genes observed compared to organoids.

As anticipated, the introduction of differentiation media results in the differential expression of many genes relating to stem cell maintenance and differentiation in the WNT, R-spondin, Notch, and TGF-β pathways. Subsequently, the alteration in urothelial-relevant transcriptional factor (e.g., MECOM) and functional gene expression (e.g., UPK1B) matches our expected hypotheses that a maturation process would occur in differentiated organoids. Several gene sets were enriched as well, representing broadly retinoid-related compound metabolism, hydrolase activity, and chemokine activity. Retinoic acid signaling is important for differentiation and maturation of the urothelium.^45^ The increase in cytokine and antimicrobial peptide gene expression supports the maturation of the cultures to potentially defend against pathogens.

### Estradiol downregulates cytokine expression while upregulating demannosylation pathways

Our exploration of the effect of estradiol on urothelial expression originated from attempting to understand the protective role it holds clinically to treat recurrent UTI in postmenopausal women (when administered locally).^9^ Unexpectedly, we observed a downregulation of several innate immunity genes, including CXCL8 (log_2_ fold-change = -0.63). Similarly, we observe differential decreases in the endogenous expression of seven genes which code for antimicrobial peptide genes including CXCL14 (log_2_ fold-change = -1.06), POMC (log_2_ fold-change = -1.50), RARRES2 (log_2_ fold-change = -0.86). This was further supported in gene set enrichment analysis where the antimicrobial humoral immune response mediated by antimicrobial peptide set was downregulated (Figure 5E). Previous work with primary urothelial cells has suggested estradiol supplementation and menopause status impacted expression of five AMPs (DEFB1, DEFB4B, S100A7, RNASE7, CAMP) and bacterial load and invasion.^7^ We observe minimal to negligible expression of any of the corresponding genes in our sterile organoid conditions. iPSC organoids for other tissues have been previously reported to be more immature than primary adult organoids and in vivo tissue so it is possible matured tubuloids could express these genes.

Estrogen signaling has been previously shown to modify serine-type endopeptidase inhibitor (serpin) activity, including SERPINE1.^46^ We similarly observe downregulation of several serpins, especially SERPINE1 (log_2_ fold-change = -1.91), which is involved in fibrinolysis and urokinase pathways (Figure S2F) via inhibition of the urokinase receptor PLAUR. PLAU (log_2_ fold-change = -0.60) and PLAUR (log_2_ fold-change = -0.51), which following exposure to estradiol were also downregulated, participate in self-reinforcing inflammatory cascades via CXCL8 signaling.^47^ Pharmacological activation/inhibition of this proposed cascade could confirm the role of serpin expression in the observed estradiol-mediated downregulation of inflammatory signaling.

Contrarily, we observe several differentially downregulated genes relating to glycosylation, reflected by a significant downregulation of the protein deglycosylation and protein demannosylation gene sets (Figure 5D, Figure S2G). Decreased expression of DPM3 (log_2_ fold-change = -0.58), demonstrated to play a fundamental role in the mannosylation of proteins by anchoring DPM1, further supports these relationships. Loss of function DPM3 mutations have been directly linked to disease relating to insufficient mannosylation (e.g., muscular dystrophy).^48^ UPEC strains bind via type 1 fimbria to mannosylated surface proteins, and where increased mannose levels increase UPEC attachment and virulence.^49^ Further work directly testing surface mannose levels via lectin staining and subsequent UPEC adherence could further support the hypothesis that estradiol supplementation aids in preventing recurrent UTI via modulating surface urothelial mannose levels.

### Transwell cultures mature following exposure to differentiation gradient and air-liquid interface

Under the impact of a differentiation gradient, Transwell cultures matured with differentially expression of WNT, R-Spondin, Notch, TGF-β genes, similarly to organoid differentiation (Figure 6B, Figure 7A). The ligands WNT2B, WNT7A, and WNT7B were all differentially upregulated in both the organoid and Transwell conditions. In parallel, we observed differential expression of several urothelial transcriptional factors and functional genes. The transcriptional factors PPARα and PPARδ were upregulated in both cultures. More specifically, the water transporter AQP4 was upregulated (log_2_ fold-change = 2.51), similarly to the organoid differentiation condition (log_2_ fold-change = 2.06). Additionally, the EGFR gene was upregulated in both the Transwell (log_2_ fold-change = 1.55) and organoid (log_2_ fold-change = 0.77) conditions. Further, several key genes associated with paracrine signaling, for example nitric oxide signaling, were differentially upregulated in Transwell cultures but not in organoids, including several serotonin receptors were upregulated including HTR1A (log_2_ fold-change = 8.40) and HTR1B (log_2_ fold-change = 5.23), which were poorly expressed in organoids. Notably, the nitric oxide and arginine production genes NOS1 (log_2_ fold-change = 8.01), NOS3 (log_2_ fold-change = 2.23), ASS1 (log_2_ fold-change = 2.69), and ARG2 (log_2_ fold-change = - 0.62) were differentially expressed. Each nitric oxide synthesis gene has been associated with different aspects of bladder function.^50^ NOS1 and NOS3 correlated with detrusor function and maximum flow rate, respectively. Additionally, nitric oxide plays a role in urinary tract host defense by delaying exponential expansion of UPEC, and can be even produced by probiotic bacteria as well.^51^

In total, 764 genes were differentially expressed in both conditions, with 546 genes uniquely so (Figure S6). In both conditions, the retinoic acid receptor signaling pathway gene set was upregulated (Transwell NES = 1.97, organoid NES = 2.11), while nitric oxide mediated signal transduction only was significantly upregulated in the Transwell condition (NES = 1.92). The neurotransmitter receptor activity gene set was upregulated in the Transwell condition (NES = 1.86) but just missed the significance threshold for the organoid condition with a similar enrichment (NES = 1.94, adjusted p-value = 0.052). The most differentially expressed component genes of the set in Transwells included serotonin (e.g., HTR1A), acetylcholine (e.g., CHRNA2), and glutamate (e.g., GRIA2) receptors. Of these, only the gene set for serotonin transport (NES = 1.78) crossed the significance threshold.

When cultures were exposed to an air-liquid interface for three additional days, some similar trends in differentially expressed genes occurred with the Transwell differentiation gradient cultures. 484 genes were shared between both conditions, with 315 genes uniquely to the conditions. The WNT ligand WNT7B was again upregulated (log_2_ fold-change = 1.33). The arginine and nitric oxide related genes ASS1 (log_2_ fold-change = 2.51) and NOS1 (log_2_ fold-change = 1.45) were again upregulated, plus SST (log_2_ fold-change = 2.13). Finally, the neuropeptide synthesis gene TAC1 was also upregulated in both the differentiation (log_2_ fold-change = 4.03) and ALI (log_2_ fold-change = 0.77) conditions.

Contrarily, strong and opposing signals relating to cell division were also present. For example, the regulation of mitotic nuclear division was downregulated in the differentiation gradient condition (NES = -1.97) and upregulated in the ALI condition (NES = 2.24). Additionally, some relevant barrier genes were opposingly regulated such as MUC4 (log_2_ fold-change differentiation = 3.37, ALI = -1.37) and IL11RA (log_2_ fold-change differentiation = 1.08, ALI = - 0.81). ALI exposure uniquely resulted in downregulation of several innate immunity related gene sets including peptide antigen binding (NES = -1.93), cytokine receptor activity (NES = -1.78), and antiviral innate immunity response (NES = -1.84).

Given that nitric oxide signaling gene expression levels were maintained or further upregulated in the ALI condition, it is possible that the multilayered organization or extended culture time of the Transwell culture compared to organoid cultures plays a role in the emergence of nitric oxide host defense architecture and signaling networks. The inducible NO synthase (NOS2)^51^ was not upregulated, which supports NOS1 and NOS3 upregulation is not due to pathogen challenge. Future work exploring the staining localization of nitric oxide synthase genes and measurement of NO in response to bacterial challenge would allow for improved understanding of nitric oxide pathways in our model.

## Data availability

The datasets in this work are available upon reasonable request.

## Acknowledgements

We thank Oshin Vellalara for her assistance in troubleshooting cell work and aiding in measurements. Additionally, we thank Janneke van de Wijgert for her sharing her perspectives on experimental design for evaluating hormonal treatments in organoids.

## Contributions

AJB and JMW conceived of the main research questions and experimental approach, with input from AMF. AJB, ZF, and AMF conducted the experiments. AJB, JB, and JMW designed the analysis, and AJB and JB performed the analysis. AJB illustrated and designed the figures. AJB and JMW wrote the manuscript. All authors discussed data and reviewed the manuscript.

