## Supplemental Figures and Tables for "Development of iPSC-derived urothelial organoids towards investigating the effect of hormones on host-defense to urinary tract infections"

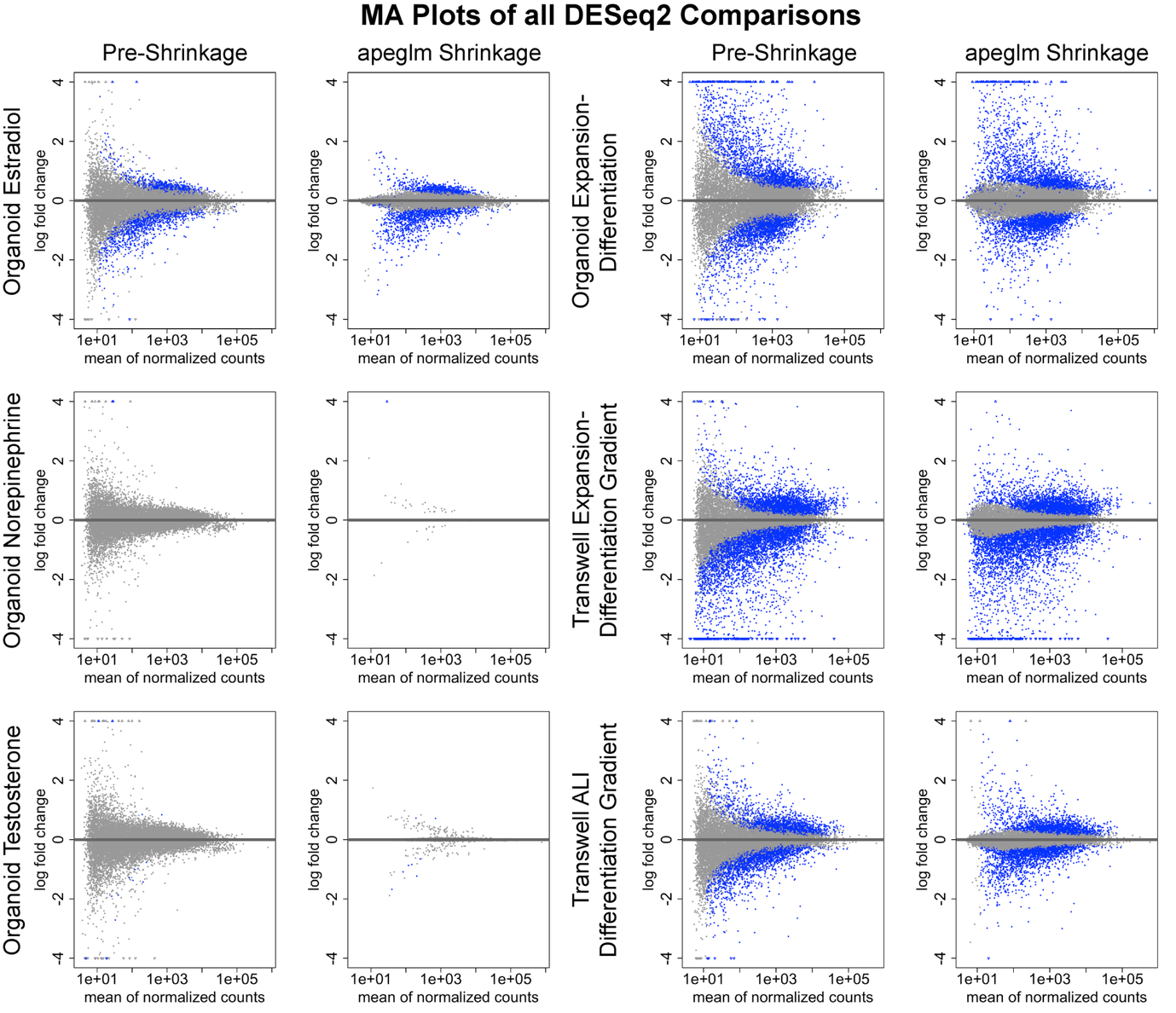

**Figure S1 –** MA plot of RNAseq comparisons before and after apeglm shrinkage plotted on a log_2_ scale for fold-change expression. Each gene represents a dot, with blue dots indicated by those below the significance threshold (adjusted *p* < 0.05).

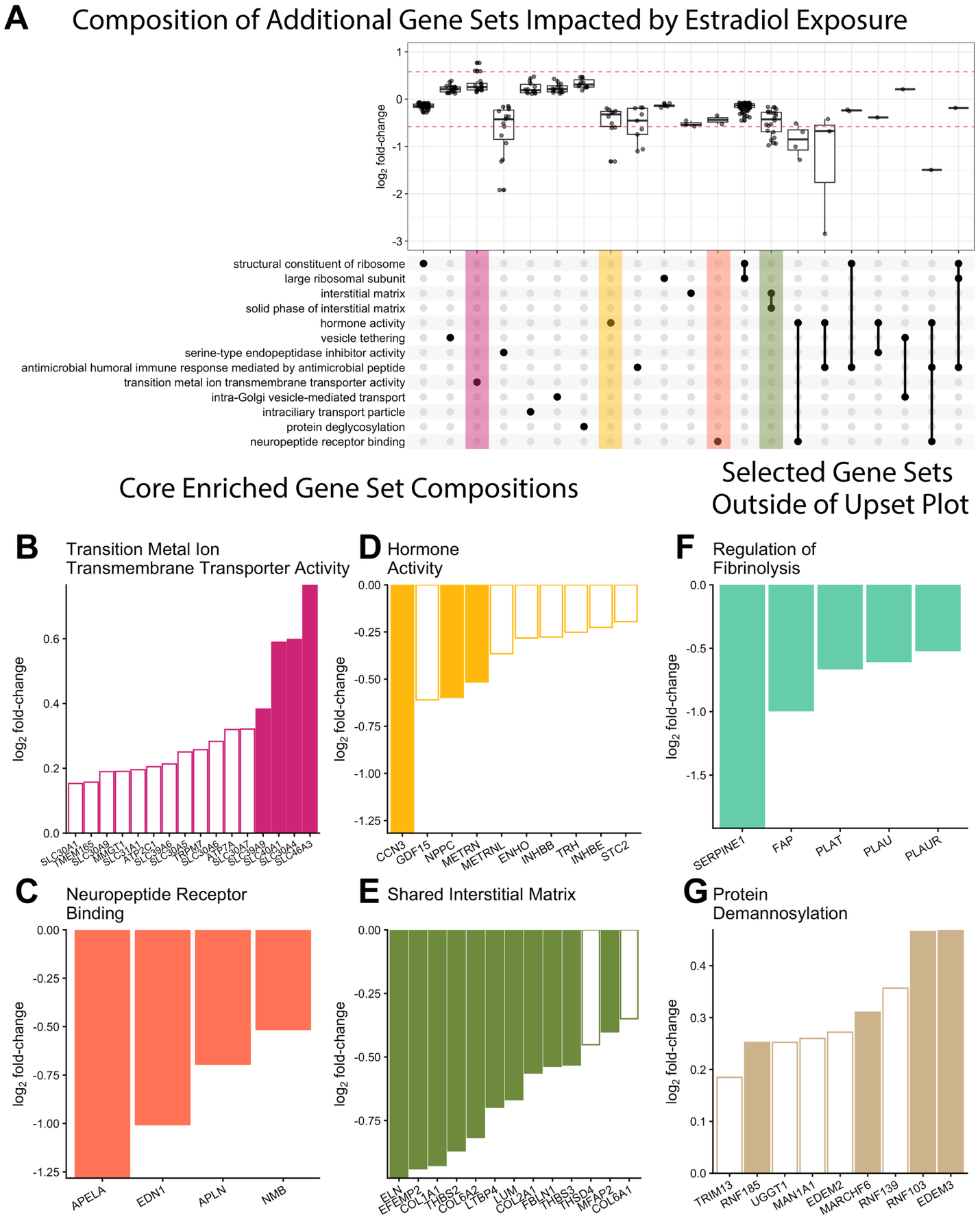

**Figure S2** – Additional enriched gene sets following estradiol treatment. A) Upset plot of the 13 significantly enriched GO terms with the highest absolute normalized enrichment scores (NES), showing shared leading-edge genes and their lod2 fold-change values. Horizontal dashed red lines indicate the log_2_ fold-change cutoff of +/- 0.58 for differential expression. leading edge genes from the B) transition metal ion transmembrane transport activity, C) hormone activity and D) neuropeptide receptor binding gene sets that were unique to each inverted gene set. E) Leading-edge genes shared between the interstitial matrix and solid phase interstitial matrix gene sets. In addition, leading edge genes from selected gene sets not included in the UpsetPlot shown in a bar chart: F) the regulation of fibrinolysis and G) protein demannosylation gene sets. Filled bars indicate genes that were significantly differentially expressed (adjusted p-value of 0.05).

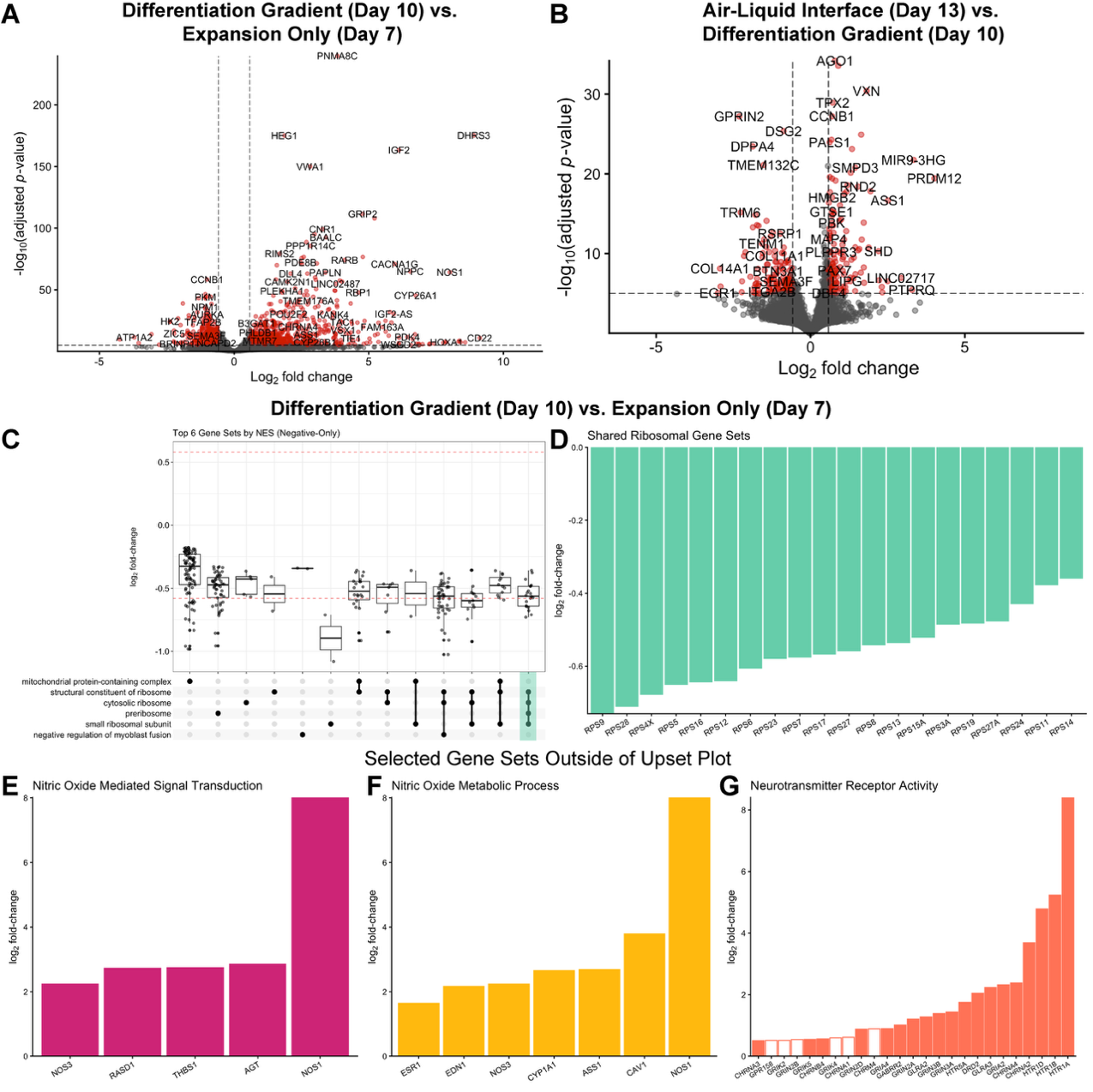

**Figure S3 –** Volcano plots of transwell group differential gene expression including the A) expansion – differentiation gradient (3317 DEGs) and B) differentiation gradient – air-liquid interface (1088 DEGs) comparisons. C) Upset plot of top six negative GO terms by normalized enrichment score from gene set enrichment analysis of the expansion – differentiation gradient comparison. Boxplots of the log_2_ fold-change expression provided per unique combination. Dotted red line indicates the 0.58 threshold used to classify genes as differentially expressed. Barplots of the log_2_ fold-change expression of leading-edge core enrichment genes within gene sets. D) Shared enriched genes between the structural constituent of ribosome, cytosolic ribosome, preribosome, and small ribosomal subunit gene sets. In addition, leading edge genes from select enriched gene sets outside the top sets by normalized enrichment score: E) nitric oxide mediated signal transduction, F) nitric oxide metabolic process, and G) neurotransmitter receptor activity. Solid filled bars indicate genes which were below an adjusted p-value of 0.05.

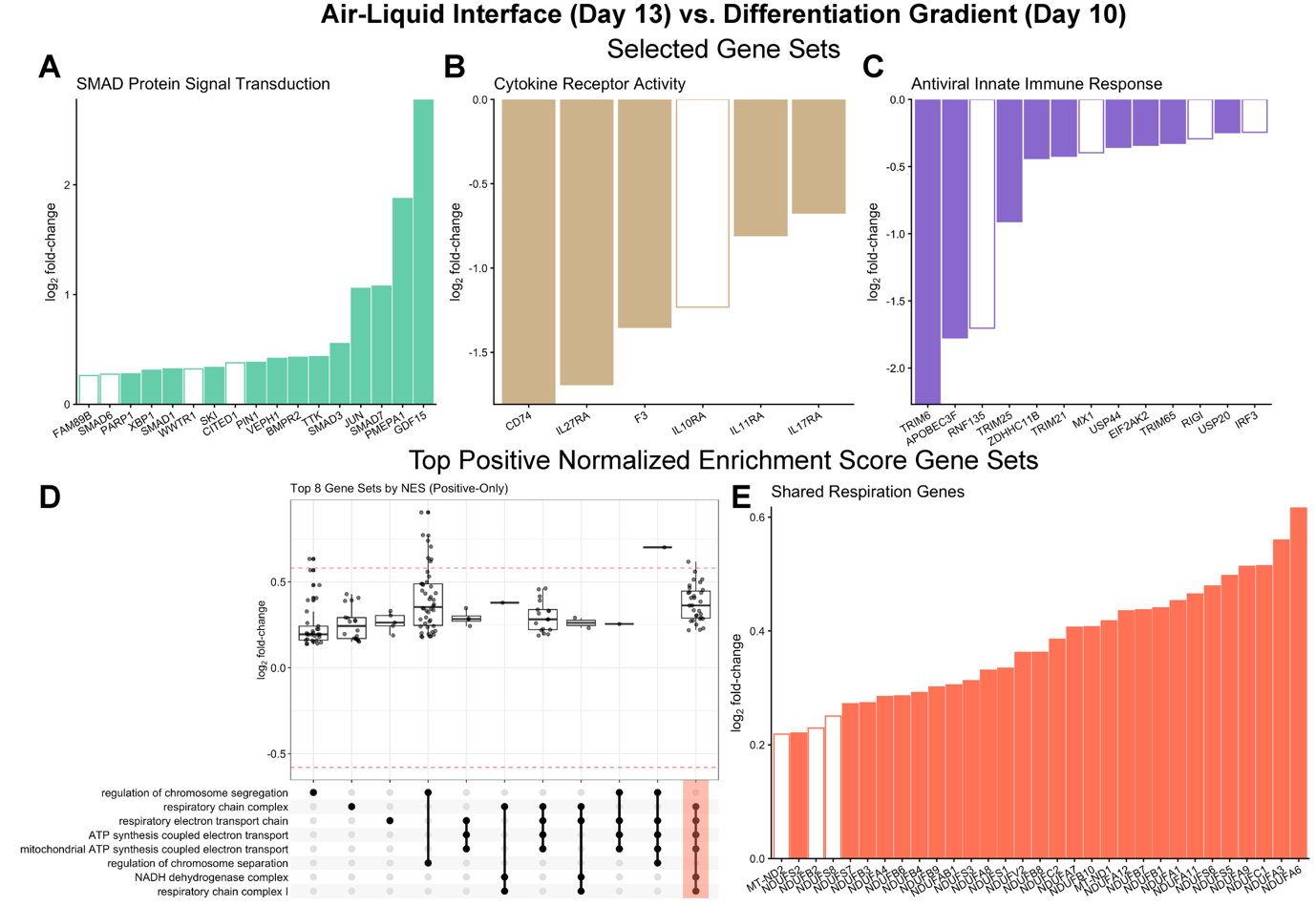

**Figure S4 –** Additional comparisons detailing the impact of an air-liquid interface on differentiation gradient urothelial barrier cultures. Enriched genes from selected significantly altered gene sets outside those included in the upset plot: A) SMAD protein signal transduction, B) cytokine receptor activity, and C) antiviral innate immune response. D) Corresponding upset plot of the top eight positive normalized enrichment score (NES) gene sets. Boxplots of the log_2_ fold-change expression provided per unique combination. Dotted red line indicates the 0.58 threshold used to classify genes as differentially expressed. E) Shared enriched genes of six of the top eight positive NES gene sets involving cellular respiration plotted on a barplot. Solid filled bars indicate genes which were below an adjusted p-value of 0.05.

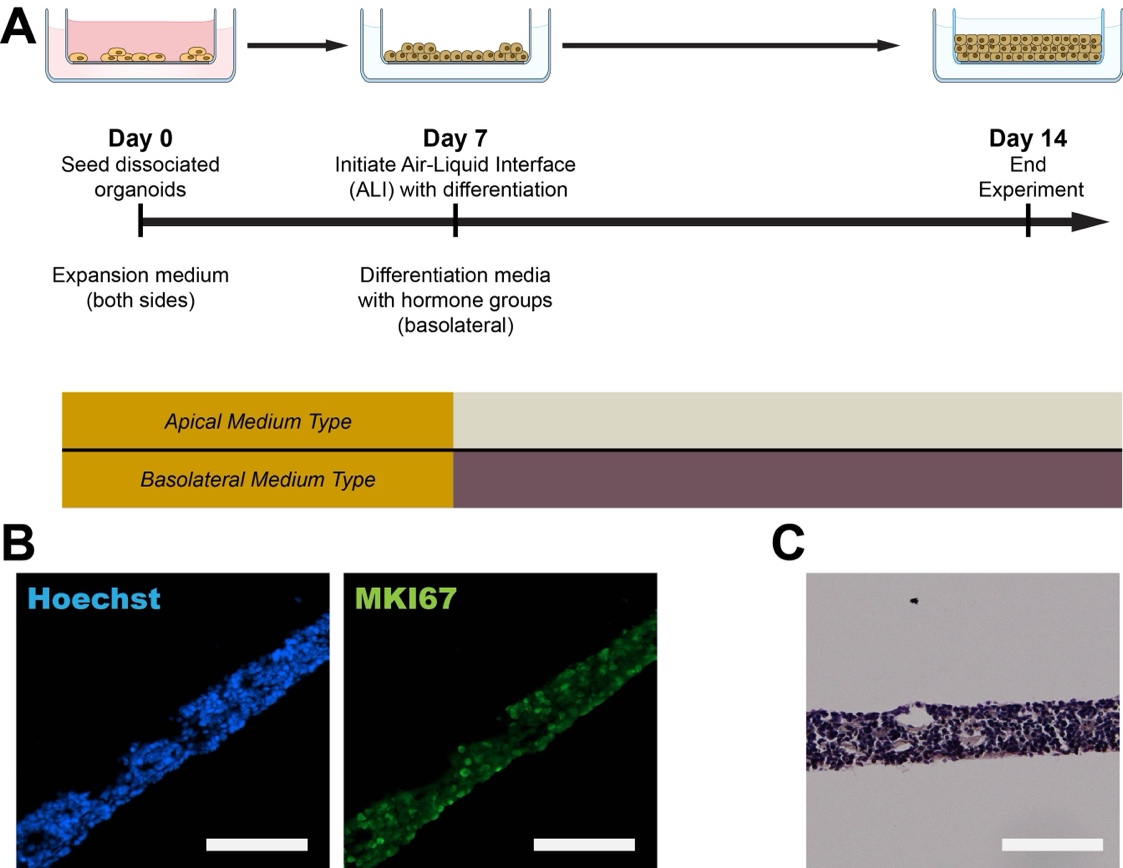

**Figure S5 –** Characterization of the multilayer barrier culture established by air-liquid interface after 14 days of culture. A) Protocol schematic. B) Immunostaining of culture slices MKI67 with Hoechst. C) H&E staining. Scale = 100 µm.

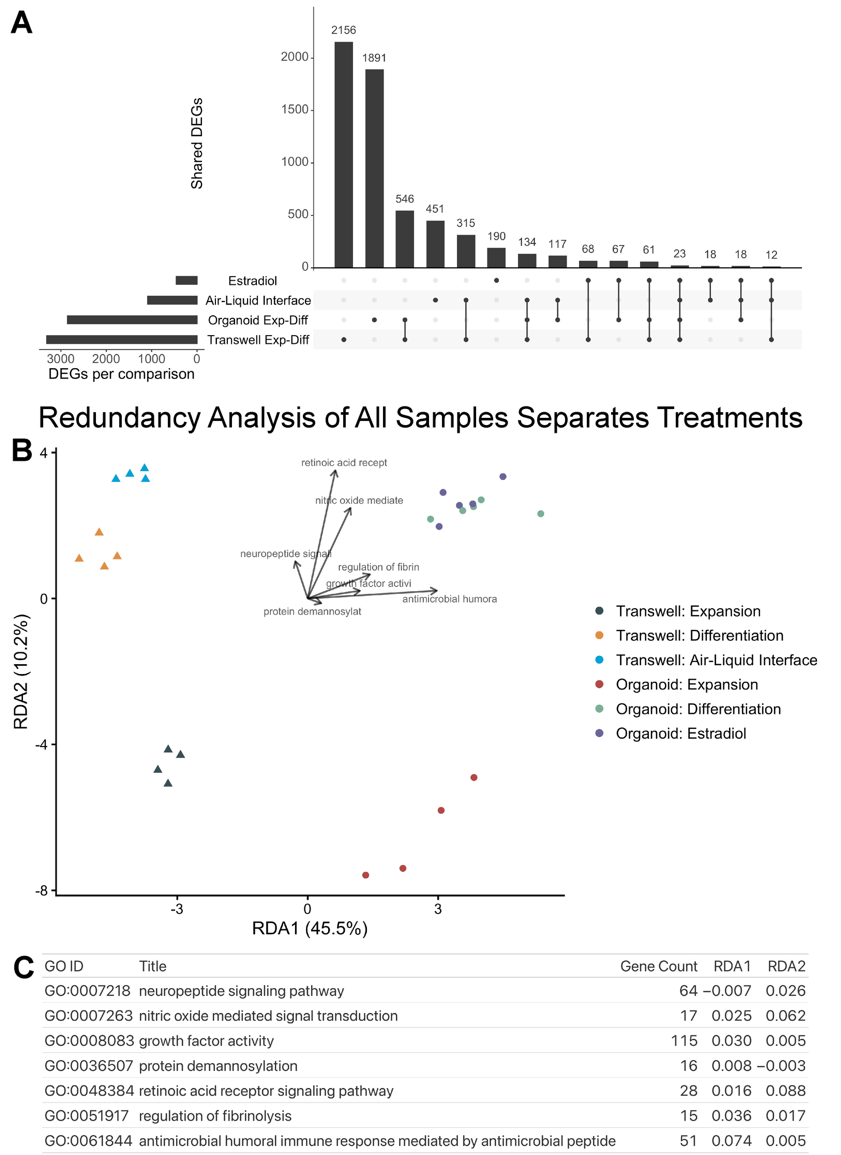

**Figure S6 –** Redundancy analysis of total samples separates strongly by group. A) Upset plot illustrating the shared DEGs between groups. B) RDA plot of all RNA-seq data with GO terms annotated. Overall model p-value < 0.001 (Table S4). Full name cutoff for legibility. C) Full information about enriched gene sets visualized in the RDA space.

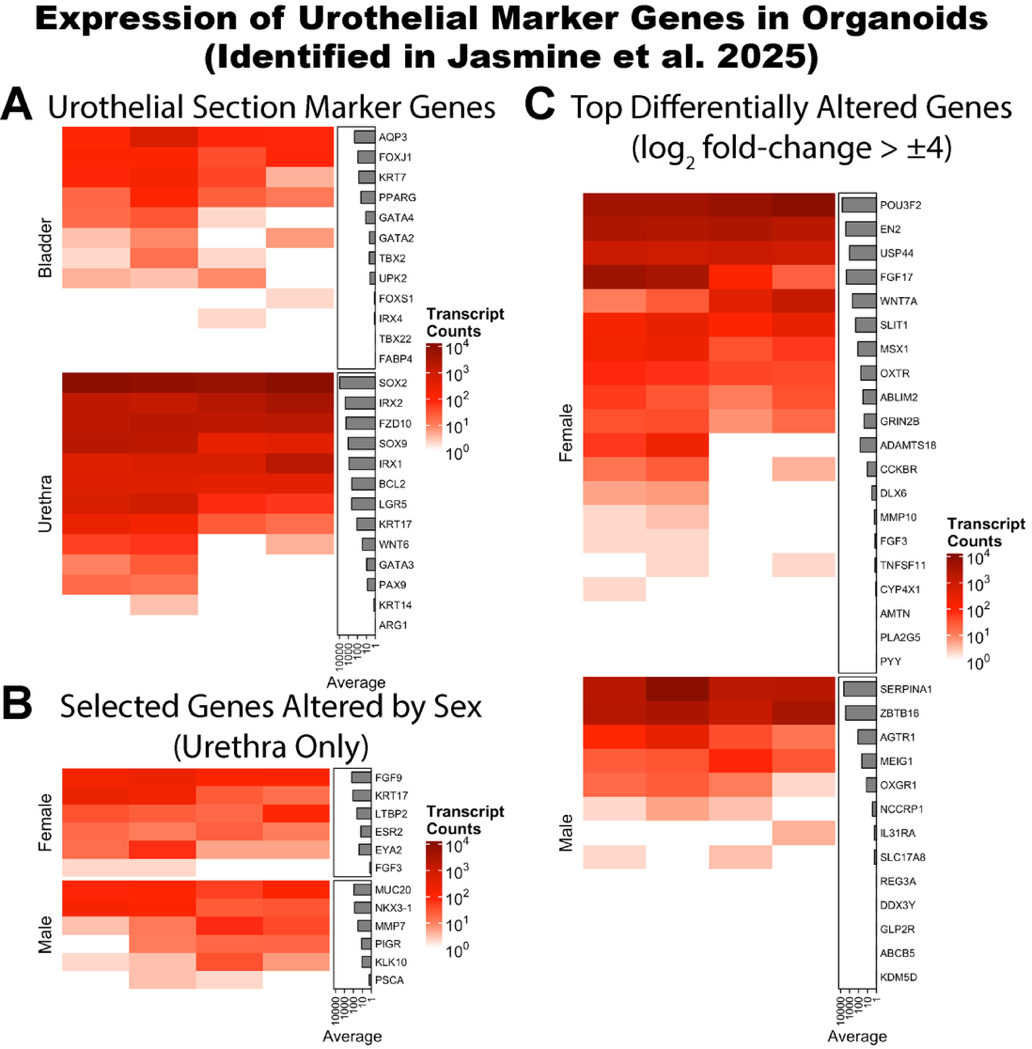

**Figure S7 –** Urothelial marker genes identified by Jasmine and Baraiya et al. 2025^40^ in mice. Included genes are human analogs identified from the Homologene NCBI database via the homologene R package. All counts are from gene-length normalized counts of organoids cultured in expansion medium where each column represents a sample. A) Urothelial genes differentially expressed by segment (bladder vs. urethra) and sex (male vs. female in urethra). B) Genes selected by Jasmine and Baraiya et al. and C) Top differentially expressed genes (log_2_ fold change > ±4).

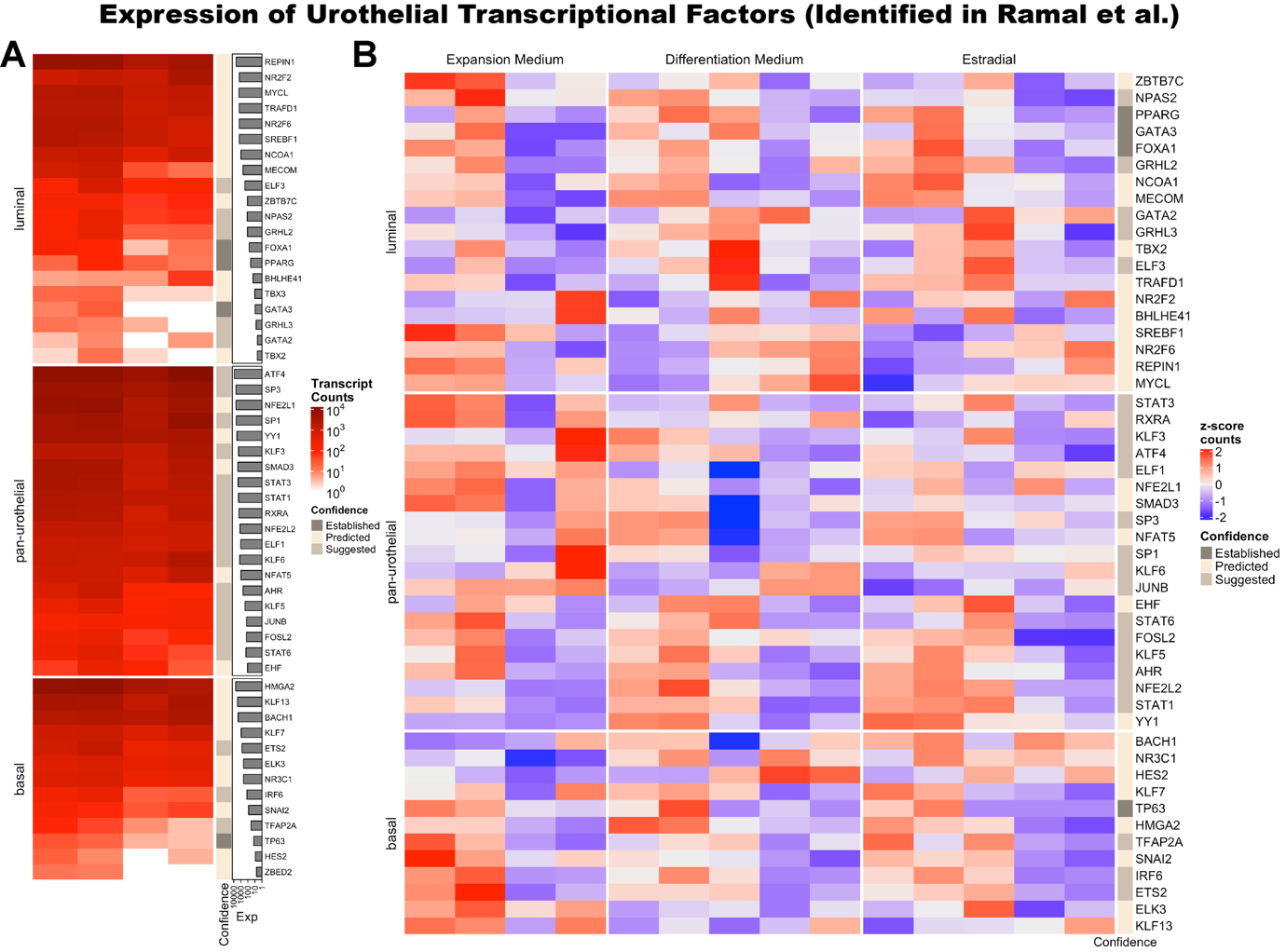

**Figure S8 –** Urothelial transcriptional factor expression in organoid cultures. Genes identified by Ramal et al.^43^ from existing and re-analyzed data and stratified by confidence level, where suggested genes were proposed in their work. A) Expression level, measured in gene-length normalized counts, of all identified genes. B) Z-score vst-normalized transcript count heatmap of organoid cultures. Each column represents a sample.

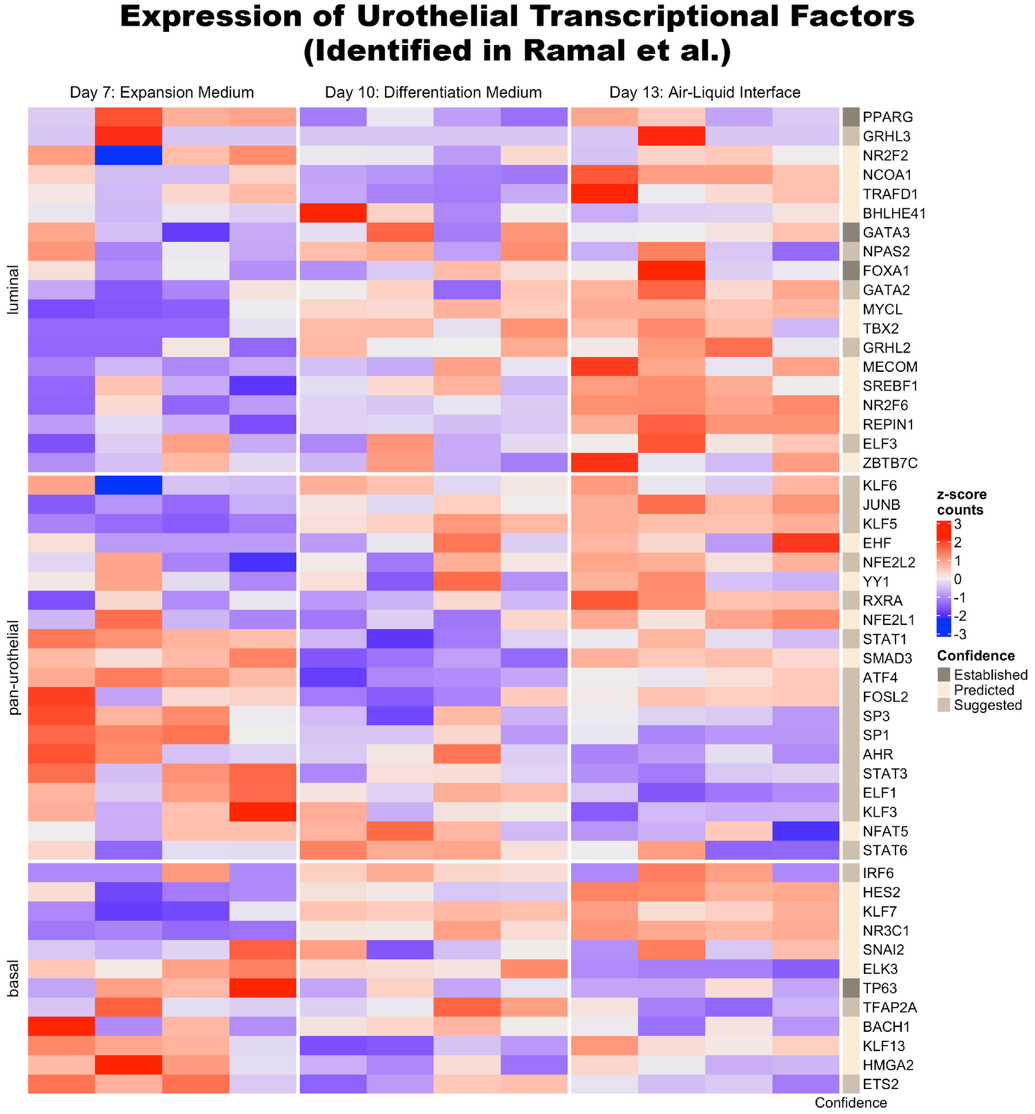

**Figure S9 –** Urothelial transcriptional factor expression in transwell cultures. Genes identified by Ramal et al.^43^ from existing and re-analyzed data and stratified by confidence level, where suggested genes were proposed in their work. Z-score vst-normalized transcript count heatmap of transwell cultures where each column represents a sample.

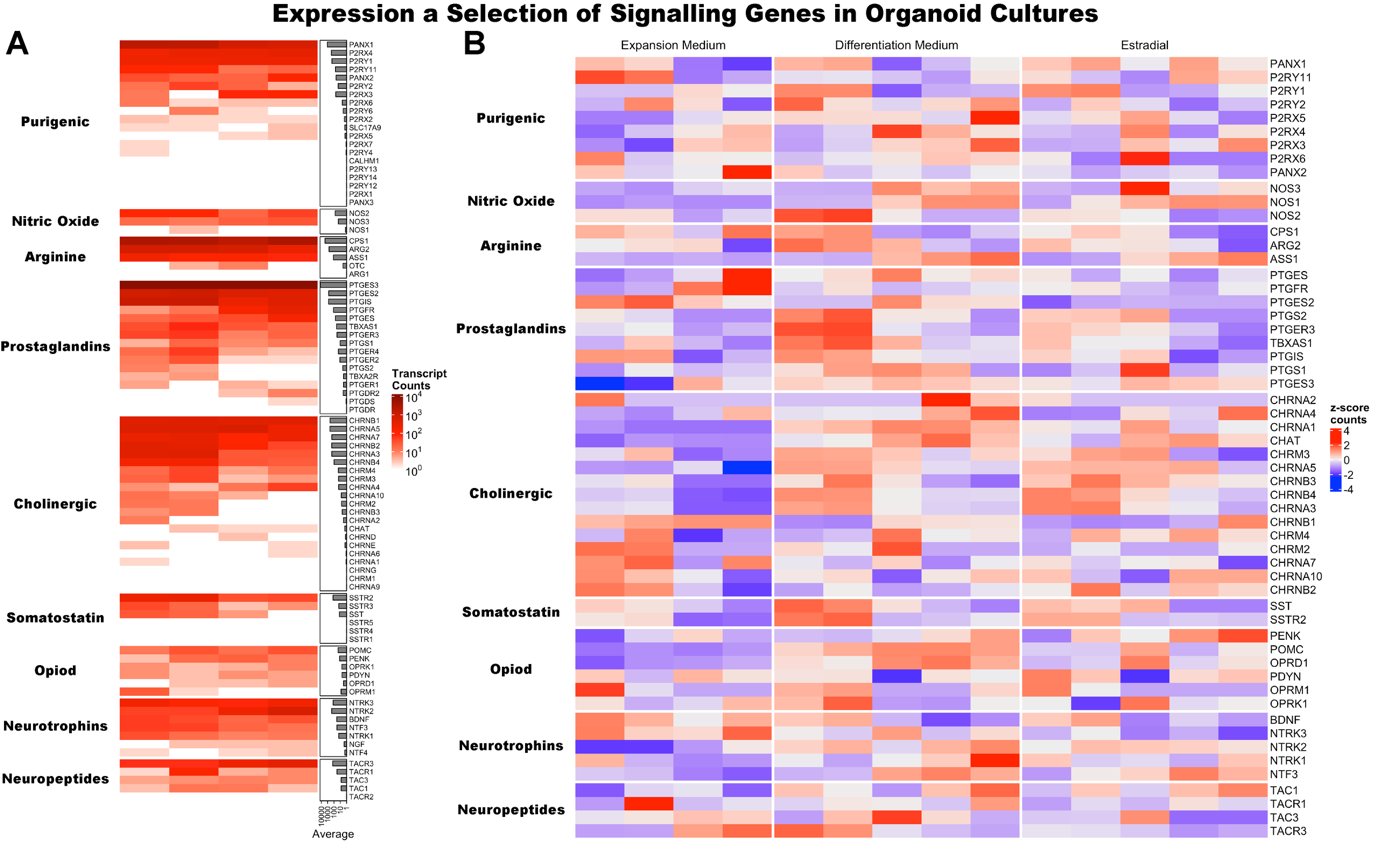

**Figure S10 –** Selected signaling pathway genes in organoid cultures. A) Expression level, measured in gene-length normalized counts, of all identified genes. B) Z-score vst-normalized transcript count heatmap of organoid cultures.

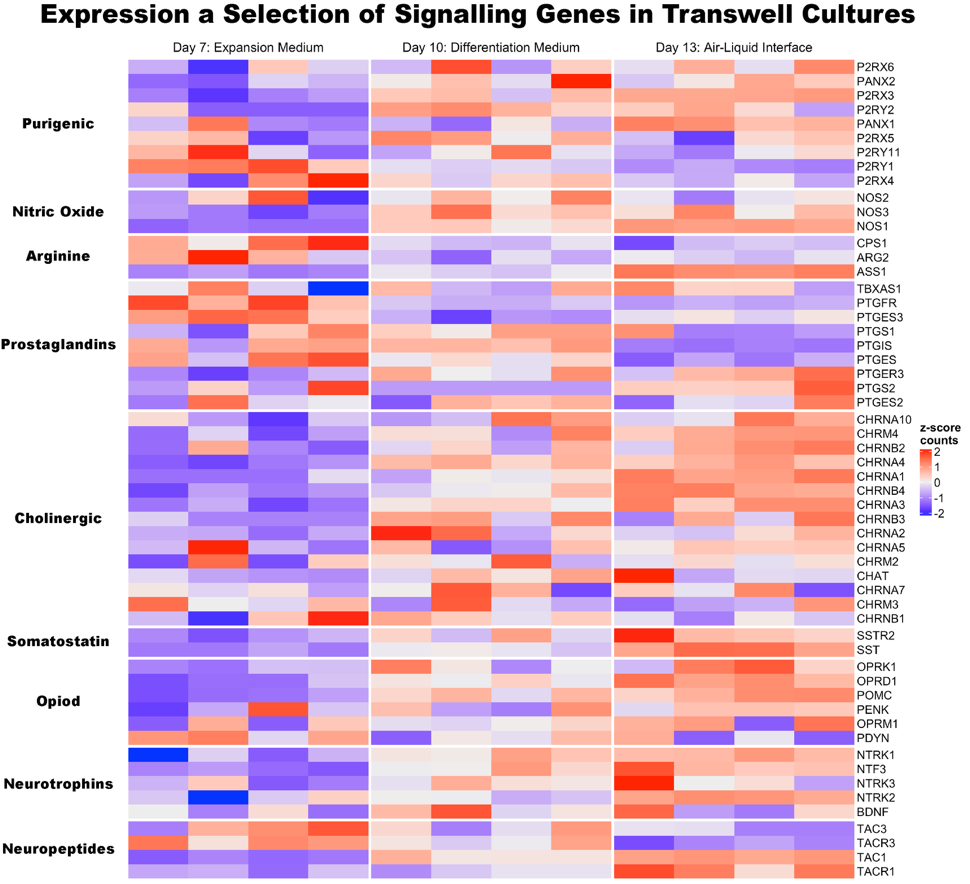

**Figure S11 –** Selected signaling pathway genes in transwell cultures. Z-score vst-normalized transcript count heatmap of transwell cultures where each column represents a sample.

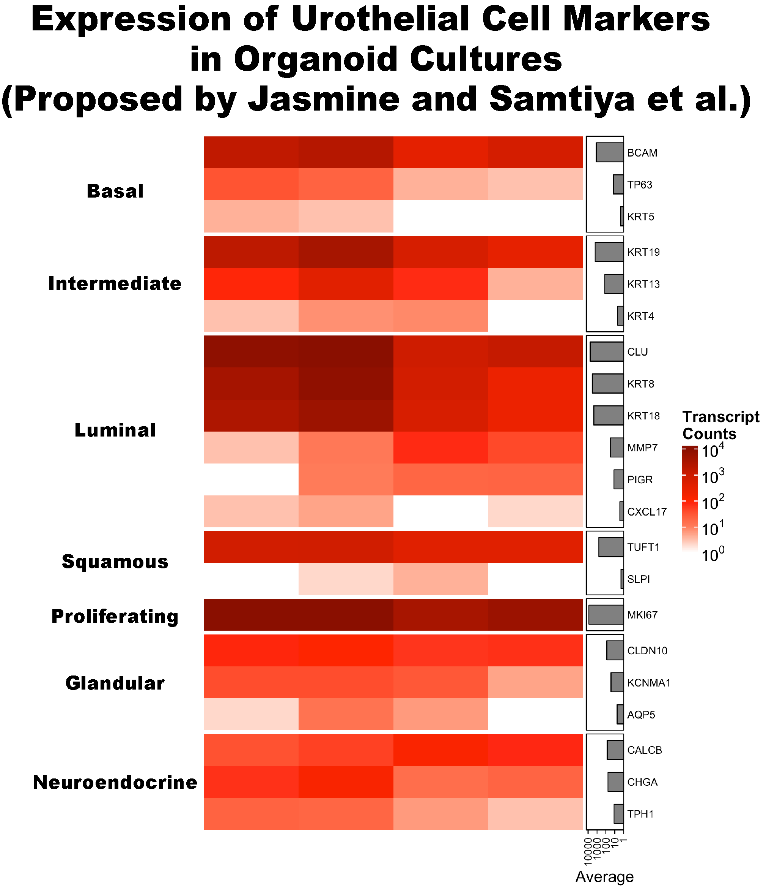

**Figure S12 –** Expression of urethral cell markers proposed by Jasmine and Samtiya et al. in organoids, measured in gene-length normalized counts where each column represents a sample.

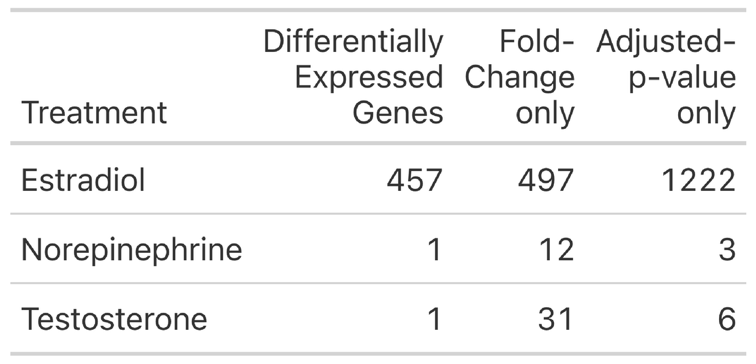

**Table S1 –** Breakdown of the impact of fold-change (absolute log_2_ fold-change > 0.58) and p-value (adjusted *p* < 0.05) together (first column) and individually only (second and third columns)

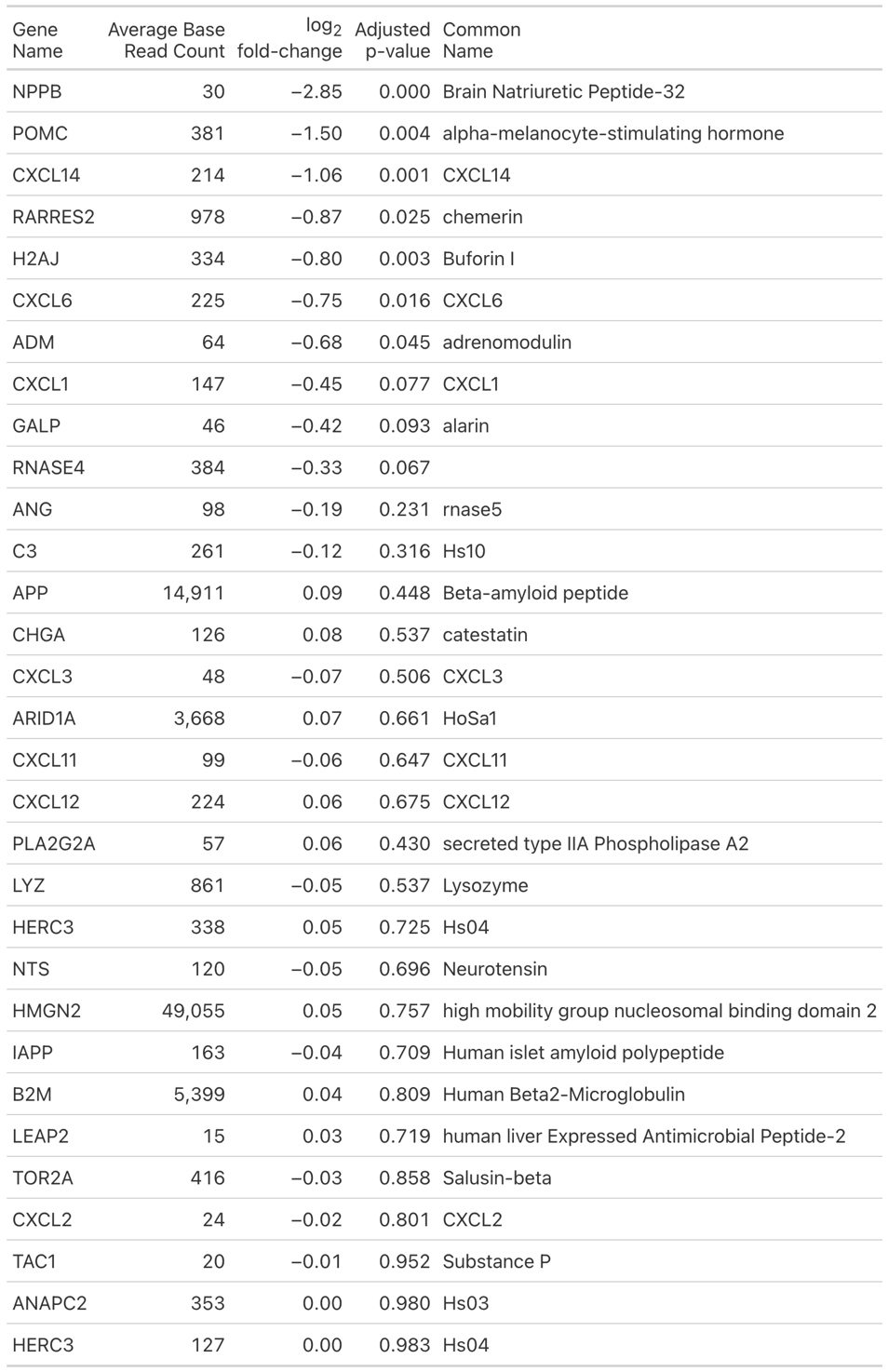

**Table S2 –** List of genes corresponding to identified antimicrobial peptides with action against gram-positive and -negative bacteria from the APD6 database, with the differential log_2_-fold expression and adjusted p-value following estradiol treatment.^37^ 99 genes were identified from the database, and 68 were due to minimal or no expression, including but not limited to: DEFB1 (hBD1), SAP1 (hBD2), S100A7, RNAS7, CAMP, DEFB4A (hBD4a), LTF (Lactoferrin), RNASE6, RNASE7.

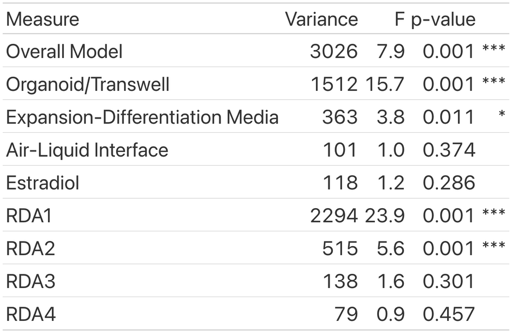

**Table S4 –** RDA model statistics calculated with 999 permutations (**p* < 0.05, ***p* < 0.01, ****p* < 0.05).

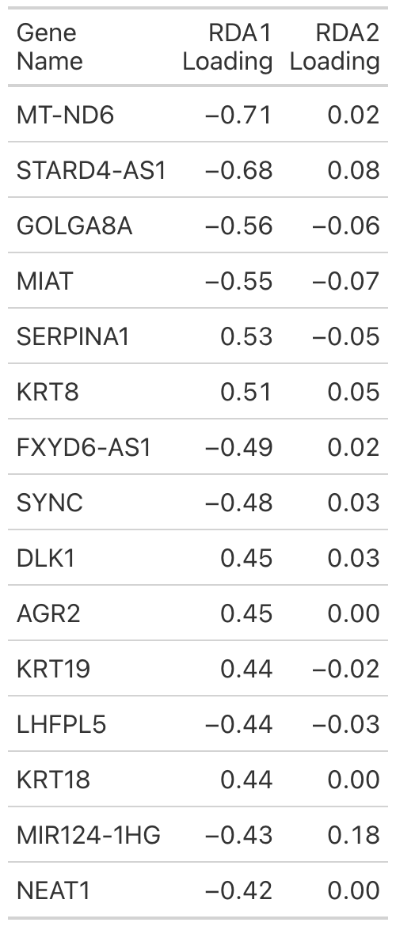

**Table S5 –** Top RDA1 loadings excluding ENSMBL IDs without a gene symbol and antisense RNA reads.

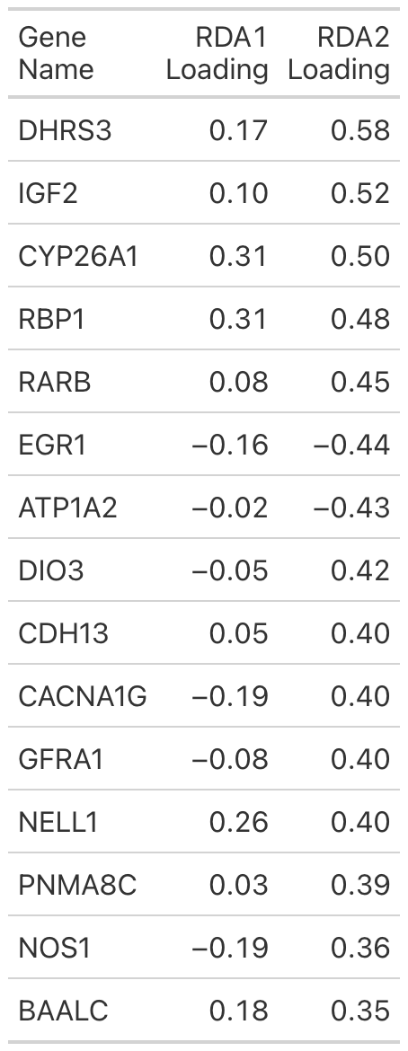

**Table S6 –** Top RDA2 loadings excluding ENSMBL IDs without a gene symbol and antisense RNA reads.
